# VE-cadherin endocytosis controls vascular integrity and patterning during development

**DOI:** 10.1101/769182

**Authors:** Cynthia M. Grimsley-Myers, Robin H. Isaacson, Chantel M. Cadwell, Jazmin Campos, Marina S. Hernandes, Kenneth R. Myers, Tadahiko Seo, William Giang, Kathy K. Griendling, Andrew P. Kowalczyk

## Abstract

Tissue morphogenesis requires dynamic intercellular contacts that are subsequently stabilized as tissues mature. The mechanisms governing these competing adhesive properties are not fully understood. Using gain- and loss-of-function approaches, we tested the role of p120-catenin (p120) and VE-cadherin (VE-cad) endocytosis in vascular development using mouse mutants that exhibit increased (VE-cad^GGG/GGG^) or decreased (VE-cad^DEE/DEE^) internalization. VE-cad^GGG/GGG^ mutant mice exhibited reduced VE-cad-p120 binding, reduced VE-cad levels, microvascular hemorrhaging, and decreased survival. By contrast, VE-cad^DEE/DEE^ mutants exhibited normal vascular permeability but displayed microvascular patterning defects. Interestingly, VE-cad^DEE/DEE^ mutant mice did not require endothelial p120, demonstrating that p120 is dispensable in the context of a stabilized cadherin. *In vitro*, VE-cadDEE mutant cells displayed defects in polarization and cell migration that were rescued by uncoupling VE-cadDEE from actin. These results indicate that cadherin endocytosis coordinates cell polarity and migration cues through actin remodeling. Collectively, our results indicate that regulated cadherin endocytosis is essential for both dynamic cell movements and establishment of stable tissue architecture.

**Summary Statement:** This study uses mouse genetic and *in vitro* approaches to demonstrate that cadherin endocytosis is critical for the formation of blood vessels during development by promoting actin-dependent collective cell migration, whereas the inhibition of this endocytosis by p120 binding is essential for vessel stabilization.

## Introduction

Collective cell movements are a central feature of tissue patterning throughout embryonic development and are essential for wound healing in adult organisms (Friedl and Gilmour, 2009; Mayor and Etienne-Manneville, 2016). Significant advances have been made toward understanding the signaling and growth factor pathways that contribute to these processes. However, we lack a comprehensive understanding of how cells engage in adhesive intercellular contacts that are adequately dynamic to allow for collective cell movement and yet sufficiently stable to maintain tissue architecture. In the present study, we used gain- and loss-of-function mouse genetic approaches to understand how endocytosis governs cadherin dynamics to permit both angiogenic vascular remodeling and vascular cohesion during development.

Blood vessel formation is fundamental to embryonic development and organogenesis as well as numerous pathological conditions ranging from diabetes to cancer (Carmeliet and Jain, 2011; Fallah et al., 2019; Folkman, 2007). Formation of the hierarchically branched vascular network is driven largely by angiogenic sprouting of endothelial cells from preexisting vessels during early development (Chappell et al., 2011; Potente et al., 2011). Sprout formation is a complex morphogenetic process that entails the polarization and collective migration of endothelial cells coordinated with proliferation, differentiation and lumen formation (Betz et al., 2016; Geudens and Gerhardt, 2011; Schuermann et al., 2014). During sprouting, contacts between neighboring cells must remain tight to maintain cohesion. However, sprouting is also highly dynamic and involves cell intercalations and coordinated cell shape changes requiring constant remodeling of cell-cell contacts (Arima et al., 2011; Bentley et al., 2014; Jakobsson et al., 2010; Szymborska and Gerhardt, 2018).

Vascular endothelial cadherin (VE-cad) is the principal cell-cell adhesion molecule of the endothelial adherens junction (Giannotta et al., 2013; Lagendijk and Hogan, 2015). The extracellular domain of VE-cad mediates adhesion through homophilic *trans* interactions, whereas its cytoplasmic tail associates with the actin cytoskeleton, providing mechanical strength to the adhesive junction (Dejana and Vestweber, 2013; Oas et al., 2013; Shapiro and Weis, 2009). VE-cad is expressed selectively in vascular and lymphatic endothelial cells and has been implicated in multiple aspects of blood vessel formation (Abraham et al., 2009; Gaengel et al., 2012; Helker et al., 2013; Lenard et al., 2013; Sauteur et al., 2014). Mice lacking VE-cad die during mid-embryogenesis due to the disintegration of nascent vessels (Carmeliet et al., 1999; Crosby et al., 2005; Gory-Faure et al., 1999), underscoring the importance of cadherin mediated adhesion to vascular development. Importantly, VE-cadherin adhesion is likely to be dynamically regulated during angiogenesis. Computational modeling and analysis of embryoid bodies and developing mouse vessels suggested that VE-cad is dynamically regulated as endothelial cells migrate collectively during angiogenesis (Arima et al., 2011; Bentley et al., 2014; Neto et al., 2018). These studies suggest that VE-cad endocytosis might contribute to collective migration and blood vessel morphogenesis *in vivo*.

The armadillo protein p120-catenin (p120) is a central regulator of classical cadherin trafficking. p120 binds to the juxtamembrane domain (JMD) in the cadherin tail and potently inhibits cadherin endocytosis and degradation by physically masking various endocytic signals (Cadwell et al., 2016; Ishiyama et al., 2010; Nanes et al., 2012). Tissue specific knockout of p120 in mice leads to decreased cadherin levels and morphogenetic defects in a variety of tissues (Davis and Reynolds, 2006; Elia et al., 2006; Hendley et al., 2015; Kurley et al., 2012; Marciano et al., 2011; Perez-Moreno et al., 2006), including the developing vasculature (Oas et al., 2010). However, p120 also performs cadherin-independent functions in the regulation of Rho family GTPases and gene transcription (Dunach et al., 2017; Kourtidis et al., 2013). Thus, p120 appears to play cadherin dependent and independent functions. However, the relative importance of these different activities of p120 to tissue morphogenesis is poorly understood.

We previously reported that p120 binding physically masks and inhibits a highly conserved three amino acid endocytic signal (DEE) within the p120-binding domain of VE-cad. Dissociation of p120 from the cadherin tail exposes the DEE motif and allows for cadherin internalization, whereas mutation of the DEE motif to alanine residues prevents VE-cad endocytosis (Nanes et al., 2012). Structural studies of E-cadherin suggest a similar mechanism by which p120 binding could inhibit a dileucine endocytic motif (Ishiyama et al., 2010). Although the role of p120 in regulating cadherin internalization is well established in mammalian cell culture model systems, data from Drosophila models suggest that p120 binding to cadherin is dispensable for fly development (Myster et al., 2003; Pacquelet et al., 2003). In addition, a recent report suggested a role for p120 in mediating cadherin endocytosis rather than inhibiting cadherin internalization (Bulgakova and Brown, 2016). Other studies in flies suggest that dissociation of p120 from Drosophila E-cadherin leads to increased E-cadherin turnover (Iyer et al., 2019). Thus, different experimental model systems have yielded conflicting views on the requirements for p120 binding to cadherin, and no vertebrate models of a cadherin mutant deficient in either p120 binding or endocytosis have been reported.

In the present study, we used a series of gain- and loss-of-function mouse genetic approaches to directly test the role of the cadherin-p120 catenin complex and cadherin endocytosis in vertebrate development. We generated homozygous VE-cad mutants lacking the DEE endocytic signal (VE-cad^DEE/DEE^) and/or adjacent residues needed for p120 binding (VE-cad^GGG/GGG^). We found that p120 binding to the VE-cad tail is essential for vessel integrity and for vascular barrier function. However, p120 binding to VE-cad can be rendered dispensable by mutating the DEE endocytic motif to eliminate cadherin endocytosis. Similarly, the VE-cad^DEE/DEE^ mutant can rescue the embryonic lethality associated with the p120-null phenotype, demonstrating that the essential function of p120 is to bind and stabilize cadherin at the cell surface. However, we also find that cadherin endocytosis is needed for normal vessel patterning. Homozygous VE-cad^DEE/DEE^ mutants exhibit impaired angiogenesis in multiple microvascular tissue beds, and impaired migration in *ex vivo* and *in vivo* migration assays. Further, we show that VE-cad endocytosis is required for actin-dependent polarization of endothelial cells prior to collective cell movement. These findings demonstrate that p120 regulation of cadherin endocytosis is an essential mechanism that governs the plasticity of cell-cell contacts during vertebrate development, and that cadherin endocytosis is integrated with polarity cues to regulate cell migration and angiogenesis.

## Results

### Generation of VE-cad mutant alleles with disrupted p120 binding and altered endocytic rates

To determine the roles of p120 binding and VE-cad endocytosis in blood vessel development and endothelial function *in vivo*, we used the CRISPR/Cas9 system to create a series of mouse knock-in VE-cad mutants with disrupted p120 binding and altered endocytic rates, as summarized in Figure 1. These mutants allowed us to dissect the specific roles of both p120 binding and VE-cad endocytosis in a mammalian *in vivo* system using endogenous VE-cad expression levels. First, we mutated highly conserved contiguous GGG residues within the core p120-binding domain to alanine residues (designated VE-cad^GGG^) (Figure 1A,B). Mutation of these GGG residues prevents p120-binding, leading to exposure of the DEE endocytic motif and cadherin destabilization (Nanes et al., 2012). Second, we mutated the DEE residues comprising the endocytic motif (designated VE-cad^DEE^)(Figure 1A,B). Replacement of these residues with alanine residues leads to a dramatic decrease in constitutive endocytosis of VE-cad from the plasma membrane (Nanes et al., 2012). The DEE mutations also lead to reduced p120-binding, since this motif lies within the p120-binding domain (Nanes et al., 2012). Finally, in the process of making the above mutants, we generated the VE-cad^ΔJMD^ allele. This allele contains an in-frame, eleven amino acid deletion in the core p120 binding domain comprising both the DEE endocytic motif and the GGG residues (Figure 1A,B). This deletion should therefore completely abrogate both DEE-mediated endocytosis and p120 binding. Founder mice for all strains were identified by PCR/RFLP analysis and confirmed by Sanger sequencing (Figure 1C).

**Figure 1.**
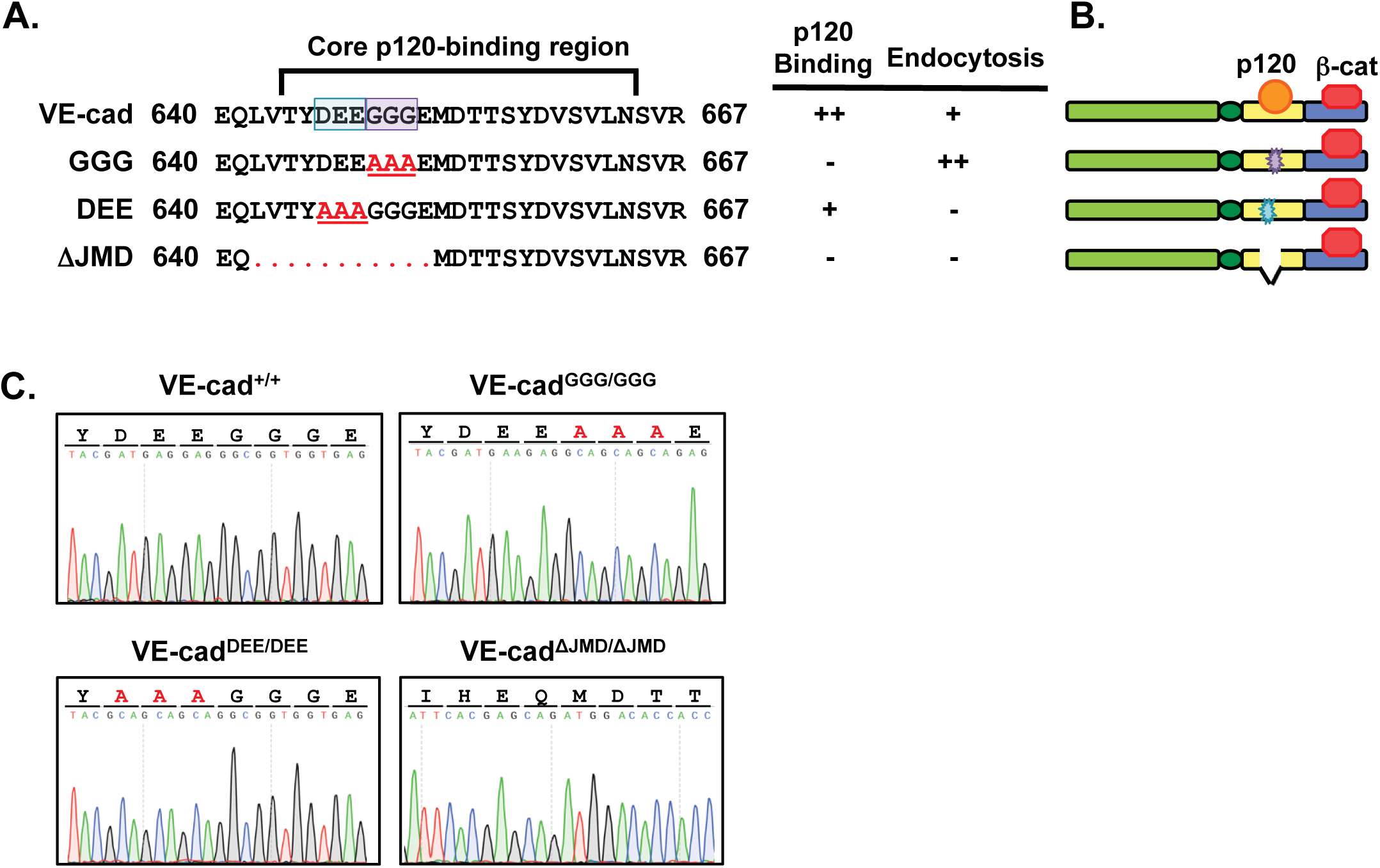
CRISPR generated VE-cad mouse mutants. **A**, Amino acid sequence of wild type VE-cad core p120-binding domain and mutants analyzed in this study. The wild type allele binds p120 and undergoes endocytosis upon p120 dissociation. The VE-cad^GGG^ allele contains alanine substitutions of GGG residues (purple box), which disrupt p120 binding leading to increased endocytosis. The VE-cad^DEE^ endocytic mutant, with mutated DEE residues (blue box), exhibits partially reduced p120 binding, yet fails to undergo endocytosis. The ΔJMD endocytic mutant contains an 11 amino deletion comprising the DEE signal and surround residues, and also fails to bind p120 or undergo endocytosis. **B**, Schematic representations of the VE-cad mutants in part A. **C**, Sanger sequencing chromatograms of the indicated VE-cadherin homozygous mutant mice.

### p120-binding to VE-cad is required for vessel integrity and survival

We first sought to determine whether p120-binding to VE-cad is required for vascular morphogenesis and endothelial function *in vivo*. Heterozygous VE-cad^GGG/+^ mice were viable, fertile, and appeared grossly normal. Although genotyping at P0 revealed homozygous VE-cad^GGG/GGG^ mutant mice were born at Mendelian ratios (25.4% out of 126 pups), these homozygous mutants often died shortly after birth. Dead VE-cad^GGG/GGG^ mutant neonates were frequently observed with visible subcutaneous spots of pooled blood, consistent with leaky blood vessels. Genotyping of offspring approximately one week after birth revealed approximately 30% of VE-cad^GGG/GGG^ mutants die during this period, indicating early postnatal lethality with partial penetrance (Figure 2A). We additionally noted that surviving VE-cad^GGG/GGG^ mutants exhibited smaller body size than wild type littermates (Figure 2B,C). To further examine the lethality phenotype, we crossed VE-cad^GGG/GGG^ mice to heterozygous mice containing a null VE-cad allele (VE-cad^STOP/+^). This allele contains an early STOP codon at amino acid 647 and thus lacks the entire β-catenin binding domain, leading to a non-functional allele (Carmeliet et al., 1999). Although heterozygous VE-cad^STOP/+^ mice are normal and viable, matings with VE-cad^GGG/GGG^ mutants yielded no viable VE-cad^GGG/STOP^ offspring (Supplemental Figure 1A). This complete embryonic lethality of VE-cad^GGG/STOP^ mice further supports a reduced function of the GGG mutant allele, and indicates that the binding of p120 is critical for VE-cad function during development.

**Figure 2.**
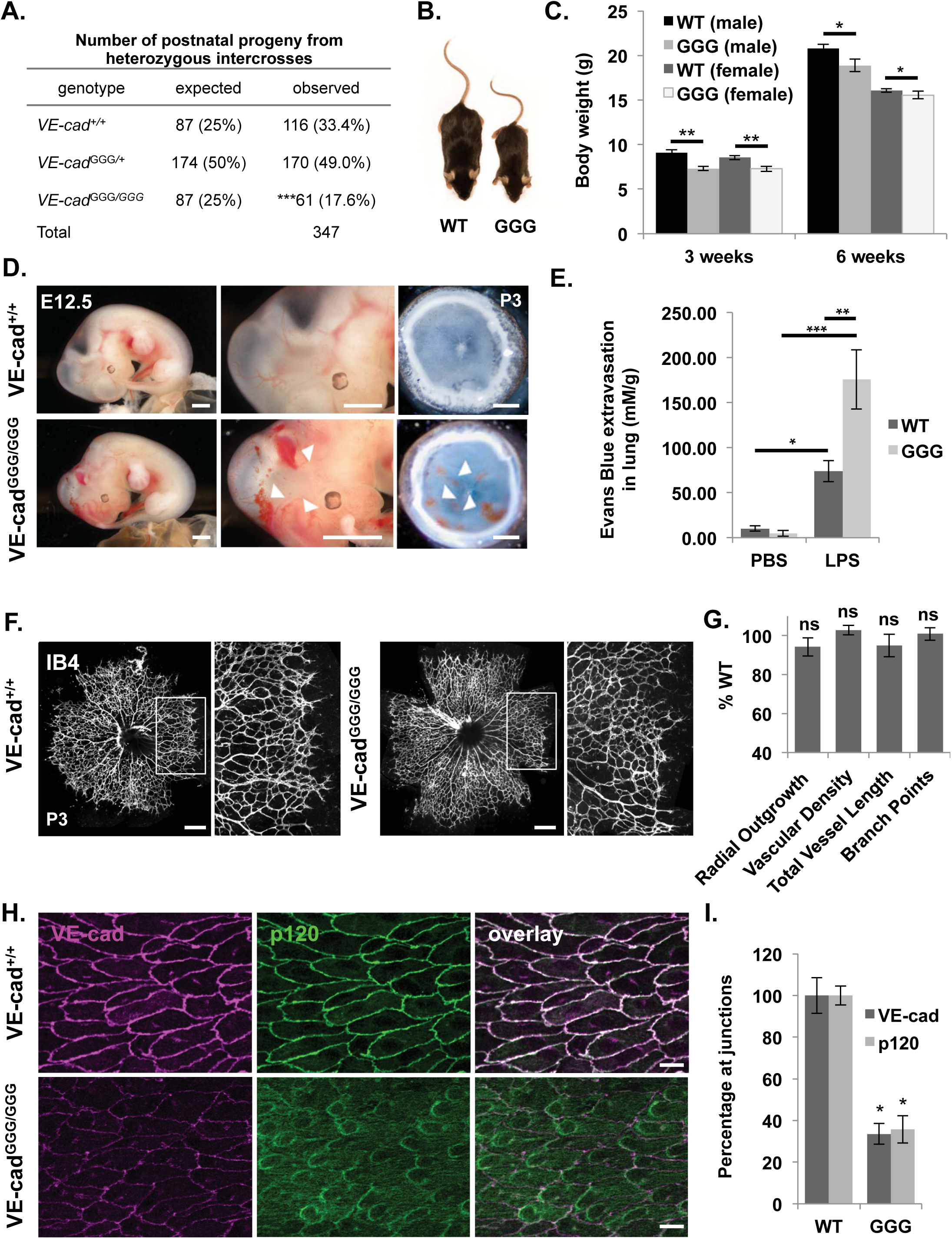
Decreased VE-cad levels, lethality and permeability defects in VE-cad GGG mutant mice lacking p120 binding. **A**, Genotyping analysis of postnatal offspring from VE-cad^GGG/+^ intercrosses reveals less than expected number of VE-cad^GGG/GGG^ homozygous mutants, indicating partial lethality. Genotyping was performed between P6-P8 and the expected number of mice was based on the total number of mice and expected Mendelian ratios. ***p<0.001 in *Χ^2^* analysis. **B**, Image of male VE-cad^+/+^ and VE-cad^GGG/GGG^ littermates at 6 weeks of age illustrating small size of VE-cad^GGG/GGG^ mutants. **C**, Decreased body weight in VE-cad GGG mutant mice at 3 weeks and 6 weeks. Body weight was assessed in VE-cad^+/+^ and VE-cad^GGG/GGG^ males and females. Results are shown as mean ± SEM. *p<0.05, **p<0.001. **D**, Left and middle panels: Gross examination of VE-cad^+/+^ (top) and VE-cad^GGG/GGG^ mutant (bottom) whole embryos at E12.5. Variable size hemorrhages (arrows) were observed in the VE-cad^GGG/GGG^ embryos, which are shown at larger magnification in the middle panels. Right panels: Fixed eye cups from VE-cad^+/+^ and VE-cad^GGG/GGG^ mutant mice at P3. Larger and more frequent hemorrhages (arrows) were observed in the retinas of VE-cad^GGG/GGG^ mutants compared to littermates. Scale bar (left and middle): 1 mm. Scale bar (right): 0.3 mm. **E**, Increased vascular permeability in VE-cad^GGG/GGG^ mutant mice. Lung permeability in three month old mice in response to LPS treatment was assessed by the Evans Blue dye method 6 hours after treatment. The lungs were harvested and dye extravasation was quantified spectrophotometrically and normalized to lung dry weight. The bar graph represents means ± SEM with 6-7 mice per group. Two-way ANOVA *p<0.0437, ***p<0.0001, **p<0.0037. **F**, Visualization of the retinal vasculature by Isolectin-B4 staining at P3 revealed normal blood vessel patterning in VE-cad^GGG/GGG^ mutant retinas (right) compared to VE-cad^+/+^ (left) littermates. Panels on the right show higher magnification of the boxed region at the vessel front in left panels. Scale bar: 300 μm. **G**, Quantitation of vascular parameters at the vessel front in VE-cad^GGG/GGG^ mutant retinas at P3. Data are presented as % of WT littermate control and represent mean ±S.E.M, n=5 independent litters. ns, not significant. **H**, Aorta en face preparations from VE-cad^+/+^ and VE-cad^GGG/GGG^ adult mice immunostained for VE-cad (red) and p120 (green). VE-cad levels at cell-cell junctions are significantly decreased in the VE-cad^GGG/GGG^ p120-binding mutant, and p120 localization shifts from the cell-cell junctions to the cytoplasm. Scale bar: 20 μm. **I**, Quantitation of VE-cad and p120 levels at cell-cell junctions in the aortas of VE-cad^+/+^ and VE-cad^GGG/GGG^ mice. Levels were quantitated from four independent experiments with 4-6 images per animal, and shown as the relative mean ± SEM. *p<0.05.

Since we observed hemorrhages in newborn VE-cad^GGG/GGG^ mutants, we analyzed the macroscopic appearance of VE-cad^GGG/GGG^ embryos at E12.5 to ascertain if hemorrhaging also occurred embryonically. Although blood vessel organization in mutants appeared grossly normal, hemorrhaging was noted in some VE-cad^GGG/GGG^ embryos (Figure 2D). The bleeding localized primarily in the head of the mutant embryos, although blood spots were also visualized in the limbs and other regions along the body wall. We also analyzed bleeding in the retina of VE-cad^GGG/GGG^ mutants at P3, a time point during early formation of the superficial vascular plexus. Although avascular at birth, the mouse retina becomes vascularized in a highly reproducible manner over the first 10 days after birth. Blood vessels form at the optic nerve at the center and then grow outward radially over the surface of the retina by sprouting angiogenesis (Fruttiger, 2007). In the retinas of VE-cad^GGG/GGG^ mutants, we observed increased multifocal bleeding (Figure 2D). These blood spots were primarily localized around the growing vascular front of the plexus, where vessels are newly formed and less stable compared to those towards the center of the plexus. Quantitation revealed a five-fold increase in blood spot area in the retinas of VE-cad^GGG/GGG^ mutants compared to wild type littermates (Supplemental Figure 1B). The blood leakage and partial lethality in VE-cad^GGG/GGG^ mutants was suggestive of an underlying endothelial integrity defect. Therefore, we assessed vascular permeability in surviving VE-cad^GGG/GGG^ mutants in response to lipopolysaccharide (LPS) stimulation using the Evans blue method (Radu and Chernoff, 2013). Under control conditions, we observed no significant differences in permeability between wild type and VE-cad^GGG/GGG^ mutants. In response to LPS, however, extravasation of Evans blue dye in the lungs was significantly higher in VE-cad^GGG/GGG^ mutants than in wild type mice (74 mM/g versus 176 mM/g; p<0.005) (Figure 2E). These data, together with the presence of hemorrhaging, indicate an essential role for p120 binding in the establishment and maintenance of endothelial barrier function and the resistance to induced vascular leak.

To investigate whether loss of p120 binding in VE-cad^GGG/GGG^ mutants may be associated with decreased vessel sprouting and branching, we assessed vascular development in VE-cad^GGG/GGG^ mutants in the postnatal retina by whole mount staining with isolectin B4. Surprisingly, we observed no gross vascular patterning changes (Figure 2F). Quantitation revealed no significant differences in radial outgrowth, vascular density, vessel length or branching between VE-cad^GGG/GGG^ mutants and wild type littermates (Figure 2G). Thus, the formation of vessels appeared normal in VE-cad^GGG/GGG^ mutants despite hemorrhage and leak, indicating that p120 binding to VE-cad is not essential for vascular patterning.

To verify that p120 binding to VE-cad was disrupted by the GGG mutation, we examined VE-cad and p120 localization in endothelial cells in WT and VE-cad^GGG/GGG^ mutants by immunostaining *en face* preparations of adult aorta. In wild type mice, we observed intense VE-cad and p120 border staining at endothelial cell junctions (Figure 2H). In contrast, p120 was absent at cell-cell borders in the aorta of VE-cad^GGG/GGG^ mice and VE-cad levels at cell junctions were dramatically reduced (Figure 2H,I). Western blot analysis of dermal endothelial cells isolated from VE-cad^GGG/GGG^ mutants revealed decreased VE-cad levels but no changes in p120 (Supplementary Figure 1C,D). There was also a similar (69%) decrease in the levels of β-catenin (β-cat) at cell borders in the mutants, consistent with the decrease in VE-cad levels (Supplementary Figure 1E,F). Since N-cadherin (N-cad) has been shown to partially compensate for VE-cad in certain contexts (Gentil-dit-Maurin et al., 2010; Giampietro et al., 2012), we also immunostained for N-cad in VE-cad^GGG/GGG^ mutant aortas, but failed to observe any upregulation of N-cad at cell borders (data not shown). Collectively, these data indicate that the loss of p120 binding to the VE-cad cytoplasmic domain leads to severe reductions in VE-cad levels and compromised endothelial barrier function, but no obvious vascular patterning defects.

### Deletion of VE-cad DEE endocytic motif restores VE-cad levels and endothelial integrity in the absence of p120-binding

We hypothesized that VE-cad^GGG/GGG^ mutants display decreased levels of VE-cad at junctions due to the inability of p120 to bind VE-cad and block endocytosis. One way to test this hypothesis would be to simultaneously disrupt both p120 binding and DEE-mediated endocytosis and determine if VE-cad levels are restored. During the process of engineering our mouse strains, we generated a mutant with an in-frame, 11-amino acid deletion encompassing both the DEE residues and GGG residues (VE-cad^ΔJMD/ΔJMD^) (Fig 1A-C). We hypothesized that this deletion should prevent both p120 binding and DEE-mediated endocytosis. In the aorta of VE-cad^ΔJMD/ΔJMD^ mutants, p120 exhibited a cytoplasmic/perinuclear pattern in endothelial cells, reminiscent of VE-cad^GGG/GGG^ mutants and consistent with the inability of the VE-cad^ΔJMD/ΔJMD^ mutant to bind p120 (Figure 3A,B). However, in contrast to the VE-cad^GGG/GGG^ mutant, VE-cad levels at endothelial cell-cell junctions in VE-cad^ΔJMD/ΔJMD^ mutants were normal (Figure 3A,B) despite the lack of p120 binding. Similarly, we observed no change in β-cat levels at cell borders in VE-cad^ΔJMD/ΔJMD^ mutants (Figure 3A,B). Total cellular levels of VE-cad and p120 were also normal in Western blots of VE-cad^ΔJMD/ΔJMD^ mutant cell lysates (Supplementary Figure 1C,D). Thus, deletion of the DEE endocytic motif prevents the downregulation of VE-cad at cell junctions associated with lack of p120 binding. These data confirm the dual function of the cadherin juxtamembrane domain in p120 binding and endocytic control. Consistent with this interpretation, VE-cad^ΔJMD/ΔJMD^ mutants appeared grossly normal and lacked the partial lethality and hemorrhaging observed in VE-cad^GGG/GGG^ mutants (Figure 3C,D and Supplementary Figure 2A). Together, these data indicate that p120 binding to VE-cad is required for normal vessel development and function, but can be rendered dispensable in the context of a VE-cad mutant that is resistant to endocytosis.

**Figure 3.**
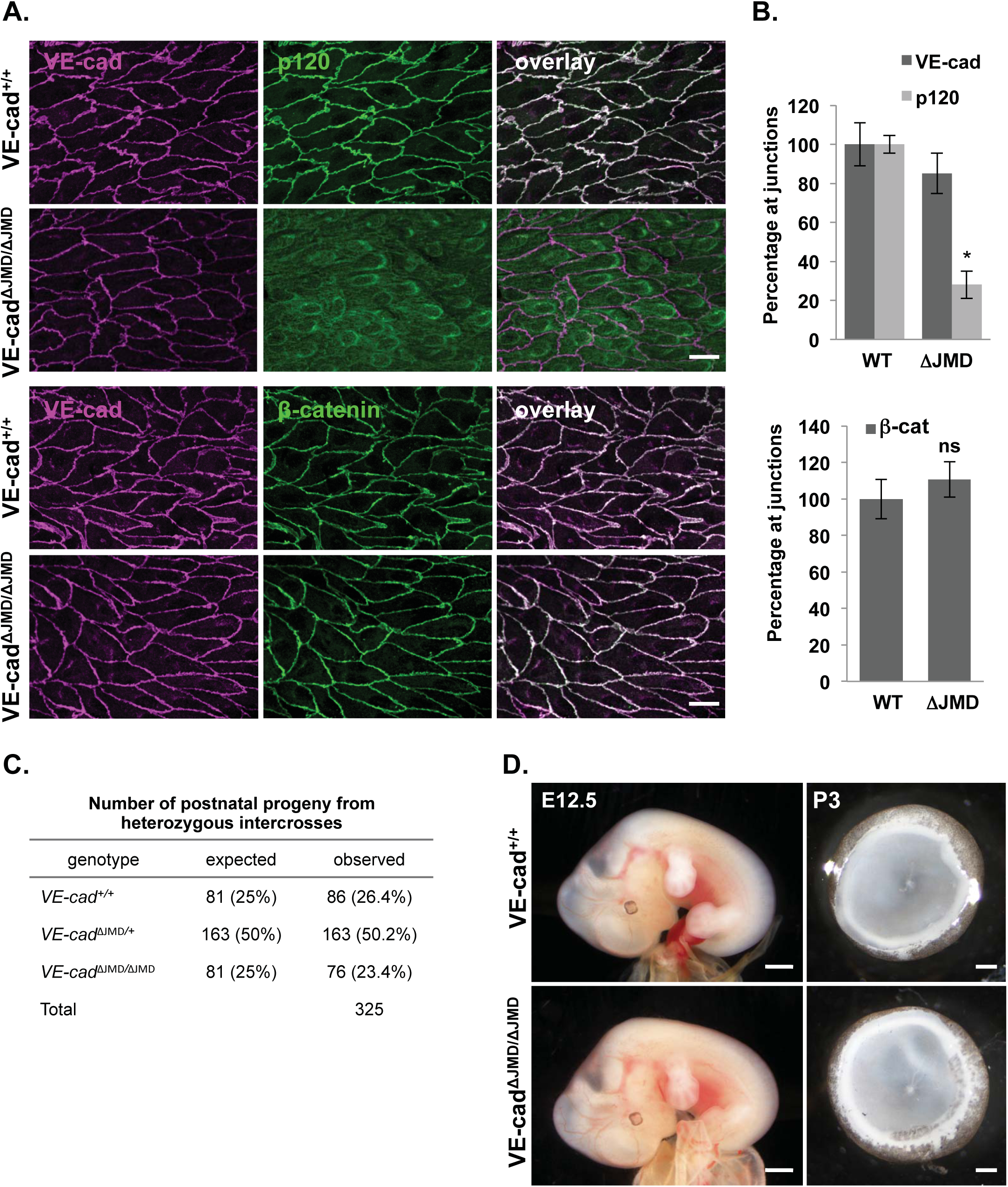
Rescue of VE-cad levels, lethality and permeability defects in VE-cad ΔJMD endocytic deletion mutant mice. **A**, Normal levels of VE-cad at cell borders in VE-cad ^ΔJMD**/**ΔJMD^ endocytic mutant mice, despite lack of p120 binding. Aorta en face preparations from VE-cad ^+/+^ and VE-cad ^ΔJMD**/**ΔJMD^ mice were immunostained for VE-cad (red) and p120 (green, top panels) or VE-cad (red) and β-catenin (green, bottom panels) and analyzed by confocal microscopy. Normal β-catenin at cell borders was also observed in VE-cad ^ΔJMD**/**ΔJMD^ mutants, corresponding to the normal VE-cad levels. Scale bar: 20 μm. **B**, Quantitation of protein levels at cell-cell junctions in the aortas of VE-cad^+/+^ and VE-cad ^ΔJMD**/**ΔJMD^ mice. No significant difference was detected in VE-cad or β-catenin levels between VE-cad ^+/+^ and VE-cad ^ΔJMD**/**ΔJMD^ mice, whereas p120 was significantly decreased at cell junctions. Levels were quantitated from four independent experiments with 4-6 images per animal, and represent the relative mean ± SEM. *p<0.05 compared to VE-cad ^+/+^; ns, not significant. **C**, VE-cad ^ΔJMD**/**ΔJMD^ mice from VE-cad ^ΔJMD /+^ intercrosses were born at normal Mendelian ratios and displayed no defects in postnatal survival. The expected number of mice was calculated based on Mendelian genetics. p=0.24 in *Χ^2^* analysis. **D**, No increase in hemorrhages in E12.5 whole embryos (left) or P3 eye cups (right) in VE-cad ^ΔJMD**/**ΔJMD^ mutant mice compared to VE-cad ^+/+^ controls. Scale bar (left): 1 mm Scale bar (right): 0.2 mm.

### Mutation of DEE endocytic motif partially rescues the p120 knockout phenotype

The normal growth and development of the VE-cad^ΔJMD/ΔJMD^ mutants indicate that p120 binding to VE-cad is not required for survival and normal vessel development. Although p120 is not bound to VE-cad in the VE-cad^ΔJMD/ΔJMD^ mutants, p120 is still present in the endothelial cells and could carry out other functions independent of cadherin binding. In previous studies, we showed that deletion of endothelial p120 resulted in embryonic lethality associated with decreased VE-cad levels (Oas et al., 2010). Here, we sought to determine if expression of an experimentally stabilized VE-cad could rescue the p120 null phenotype.

To test this possibility, we generated and characterized homozygous VE-cad mutant mice with DEE to AAA substitutions (VE-cad^DEE/DEE^) (Figure 1A-C). As discussed above, this mutation impairs VE-cad endocytosis in cultured cells (Nanes et al., 2012). We observed significantly decreased levels of p120 at cell borders in the aorta endothelium in these VE-cad^DEE/DEE^ mutants (Figure 4A,B) consistent with weak binding to p120 (Nanes et al., 2012). Nevertheless, VE-cad levels were normal in VE-cad^DEE/DEE^ mutants, with no visible difference compared to wild type mice (Figure 4A,B). β-cat levels at cell junctions were also normal in VE-cad^DEE/DEE^ mutants (Supplementary Figure 2B,C). No changes in VE-cad or p120 levels were observed in western blots of dermal endothelial lysates from these mice (Supplementary Figure 1C,D), and we observed no upregulation of N-cad in the aorta endothelium (data not shown). Furthermore, we observed expected Mendelian ratios of homozygous VE-cad^DEE/DEE^ mutants one week after birth (24.0%), indicating no lethality in these homozygous mutants (Figure 4C). VE-cad^DEE/DEE^ mutants also displayed no macroscopic hemorrhaging either embryonically or postnatally, and displayed no increase in lung permeability in response to LPS challenge (Figure 4D,E and Supplementary Figure 2A). Together, these data suggest that endothelial integrity and barrier function is normal in these mice, similar to VE-cad^ΔJMD/ΔJMD^ mutants.

**Figure 4.**
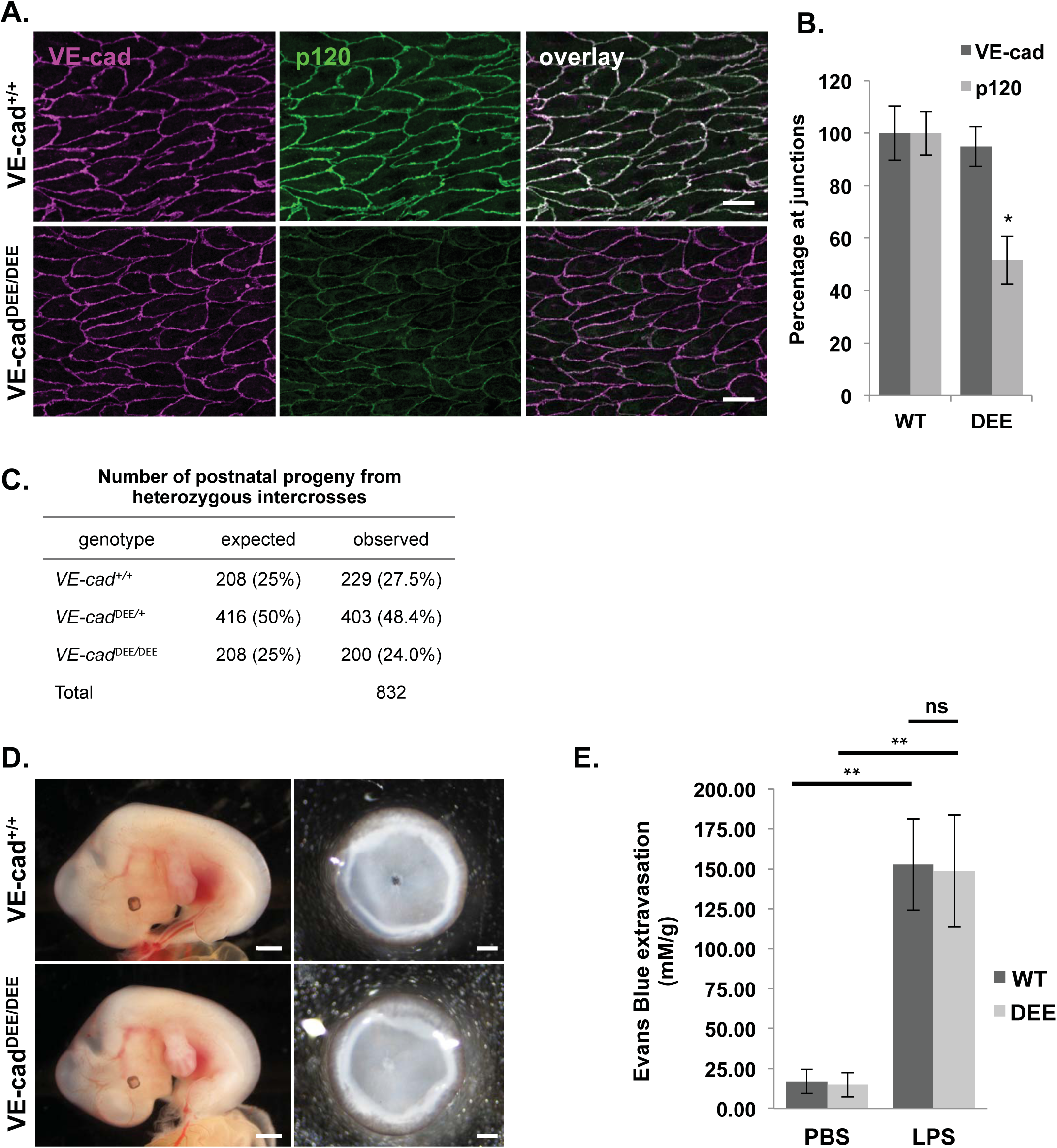
No lethality or permeability defects in DEE endocytic mutant mice. **A**, Immunostaining analysis for VE-cad (red) and p120 (green) on en face aorta preparations from VE-cad^+/+^ and VE-cad^DEE/DEE^ adult mice. VE-cad levels at cell junctions appeared normal whereas p120 levels were partially decreased. Scale bar: 20 μm. **B**, Quantitation of VE-cad and p120 levels at cell-cell junctions in the aortas of VE-cad^+/+^ and VE-cad^DEE/DEE^ mice. Levels were quantitated from five independent experiments with 4-6 images per animal, and shown as the relative mean ± SEM. *p<0.05. **C**, VE-cad ^DEE**/**DEE^ mice from VE-cad ^DEE /+^ intercrosses were born at normal Mendelian ratios and displayed no defects in postnatal survival. p=0.49 in *Χ^2^* analysis. **D**, Whole embryos at E12.5 (left) and P3 eye cups (right) from VE-cad^+/+^ and VE-cad^DEE/DEE^ mice. No intraretinal hemorrhaging was observed. Scale bar: 1 mm (left), 0.2 mm (right). **E**, No increase in vascular permeability in VE-cad^DEE/DEE^ mutant mice in response to LPS treatment. Lung permeability was assessed in adult VE-cad^+/+^ and VE-cad^DEE/DEE^ mice by the Evans blue dye method 6 hours after DPBS or LPS treatment. The bar graph represents means ± SEM with 7 mice per group. Two-way ANOVA **p<0.005; ns, not significant.

The generation of the VE-cad^DEE/DEE^ mutant mouse strain allowed us to test the possibility that the p120 null phenotype could be rescued by this stabilized cadherin mutant. We crossed VE-cad^DEE/DEE^ mutants with p120 conditional knockout mice harboring a p120 floxed allele (Davis and Reynolds, 2006) and utilized animals expressing Cre recombinase from the Tie2 promoter to delete endothelial p120. Specifically, we mated Tie2-Cre^+^; VE-cad^DEE/+^; p120^flox/flox^ males to VE-cad^DEE/+^; p120^flox/flox^ females. Based on normal expected Mendelian ratios, we expected 12.5% of the resulting offspring to be Tie2-Cre^+^; VE-cad^+/+^; p120^flox/flox^. However, we observed very few Tie2-Cre^+^; VE-cad^+/+^; p120^flox/flox^ mice (4/156 or 2.6%) (Figure 5A), consistent with our previous findings that deletion of endothelial p120 is embryonic lethal (Oas et al., 2010). Interestingly, survival was largely rescued in p120 knockout mice expressing a stabilized VE-cad, i.e., Tie2-Cre^+^; VE-cad^DEE/DEE^; p120^flox/flox^ (19/156 or 12.2%)(Figure 5A). Thus, an experimentally stabilized cadherin rescues the lethality associated with genetic deletion of p120.

**Figure 5.**
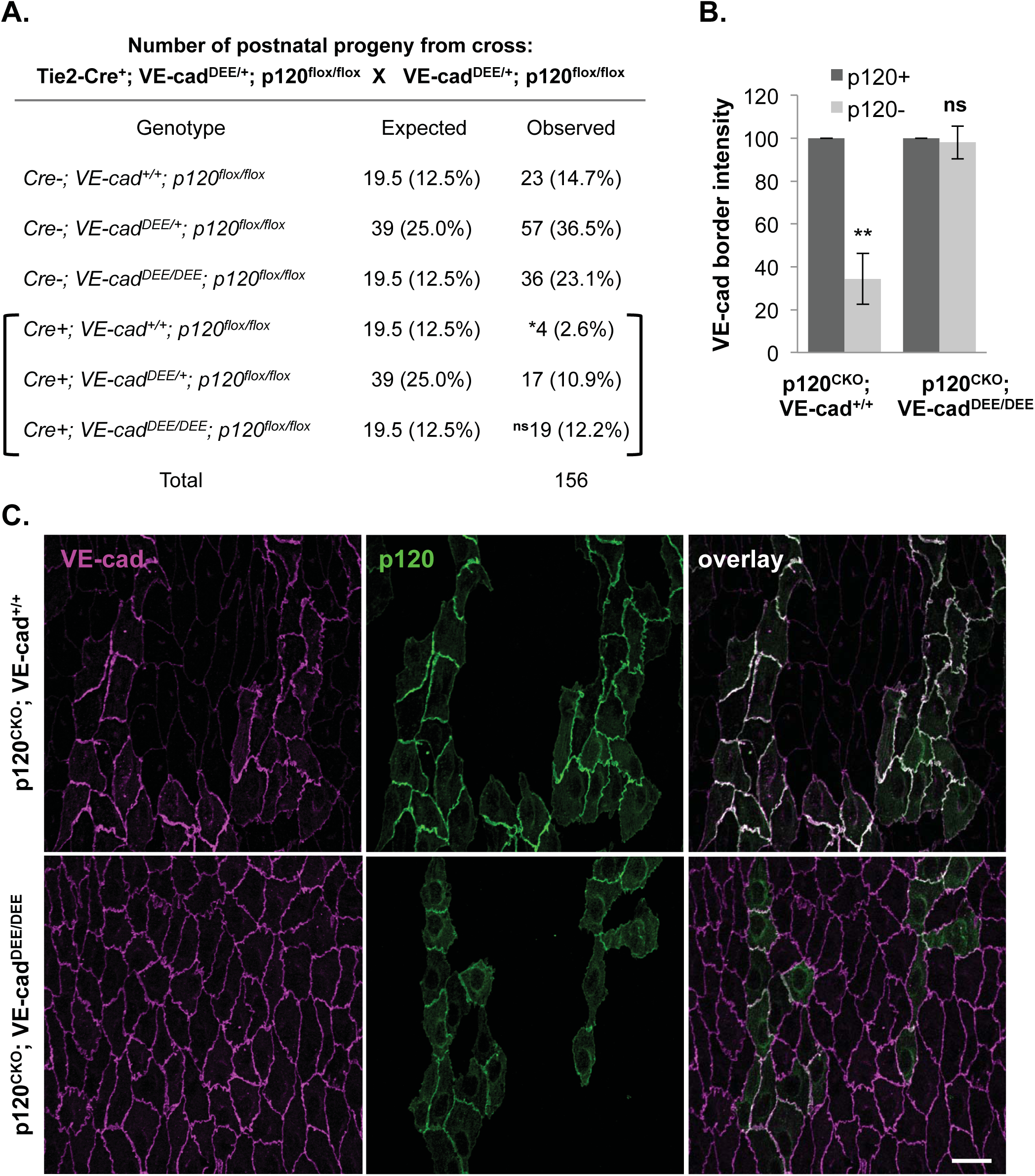
Rescue of VE-cad levels and lethality in p120 conditional knockout mice with mutation of VE-cad endocytic motif. **A**, Genotyping analysis of postnatal offspring from Tie-2-Cre^+^; VE-cad^DEE/+^; p120^fl/fl^ × VE-cad^DEE/+^; p120^fl/fl^ matings. Less than expected numbers of Tie-2-Cre^+^; VE-cad^+/+^; p120^fl/fl^ mice were born, based on expected Mendelian ratios, suggesting significant perinatal lethality in the presence of wild type VE-cad. However, Tie-2-Cre^+^; VE-cad^DEE/DEE^; p120^fl/fl^ mice were born near expected ratios, suggesting a rescue of lethality with disruption of the VE-cad endocytic motif. Genotyping was performed between P6-P8 and the expected number of mice was based on the total number of mice and Mendelian genetics. *p<0.05; ns, not significant in *Χ^2^* analysis. **B**, Quantitation of VE-cad levels at cell borders between adjacent p120^+^ or adjacent p120^−^ cells in the aortas of VE-cad-Cre^+^; p120^fl/fl^; VE-cad^+/+^ (p120^CKO^; VE-cad^+/+^) and VE-cad-Cre^+^; p120^fl/fl^; VE-cad^DEE/DEE^ (p120^CKO^; VE-cad^DEE/DEE^) mice as shown in panel C. VE-cad levels at p120^+^ cell borders were set to 100 and the percentage decrease in 120^−^ cells was quantitated. Graph represents the relative mean ± SEM, calculated from three mice per genotype **p<0.005 compared to p120^+^; ns, not significant. **C**, Rescue of VE-cad levels in p120-null cells by mutation of the DEE endocytic motif. Aorta en face immunostaining of p120^CKO^; VE-cad^+/+^ (top panels) or p120^CKO^; VE-cad^DEE/DEE^ mice (bottom panels). Mosaic Cre-mediated deletion of p120 led to both p120^+^ (green) and p120^−^ cells within the same field of view. In p120^CKO^; VE-cad^+/+^ mice, VE-cad (red) levels were significantly decreased in p120^−^ cells, suggesting p120 is required for VE-cad membrane stability. In p120^CKO^; VE-cad^DEE/DEE^ mice, no decreased in VE-cad levels were observed in p120^−^ cells, suggesting disruption of the DEE endocytic signal can stabilize VE-cad membrane levels in the absence of p120-binding. Scale bar: 25 μm.

The low birth rate of Tie2-Cre^+^; VE-cad^+/+^; p120^flox/flox^ mice precluded analysis of VE-cad levels in these mice. Therefore, we used the endothelial-specific VE-cad-Cre driver to delete p120 for further analysis. The VE-cad-Cre driver deletes p120 less efficiently than the Tie2-Cre driver, leading to a higher degree of mosaicism and a higher rate of survival of p120 conditional knockout mice. We therefore generated VE-cad-Cre^+^; VE-cad^+/+^; p120^flox/flox^ (p120^CKO^; VE-cad^+/+^) mice and VE-cad-Cre^+^; VE-cad^DEE/DEE^; p120^flox/flox^ (p120^CKO^; VE-cad^DEE/DEE^) mice and analyzed aortic endothelial VE-cad levels in p120 null (p120^−^) cells. In p120^CKO^; VE-cad^+/+^ mice, there was a dramatic decrease in VE-cad at borders between p120 null cells compared to cells expressing p120 (Figure 5B,C). However, in p120^CKO^; VE-cad^DEE/DEE^ mice, there was virtually no difference in VE-cad levels between p120^+^ and p120^−^ cells (Figure 5B,C). Collectively, these results indicate that p120 stabilizes VE-cad *in vivo* through inhibition of the DEE endocytic signal, and that p120 inhibition of VE-cad endocytosis is the primary endothelial cell function of p120 necessary for survival.

### Angiogenesis defects in DEE mutant mice

Endothelial sprouting during angiogenesis is thought to require the modulation of adherens junctions in order to permit cell intercalations and collective movements during vessel formation (Bentley et al., 2014; Szymborska and Gerhardt, 2018). Cadherin endocytosis has been implicated in the dynamic cell-cell associations required for collective migration, but this possibility has not been tested directly. We hypothesized that blood vessel formation may be altered in VE-cad^DEE/DEE^ mutants due to decreased VE-cad turnover and decreased junction plasticity. To explore this possibility, we analyzed angiogenesis in the postnatal retinas of VE-cad^DEE/DEE^ mutants. Interestingly, VE-cad^DEE/DEE^ mutants exhibited a decrease in vascular density, total length of vessels, as well as decreased vessel branching (Figure 6A,B). However, we did not observe an increase in blind-ending vessels in the mutant, indicating the angiogenesis defects were due to failures in initial endothelial sprout formation rather than vessel stabilization (Figure 6A and data not shown).

**Figure 6.**
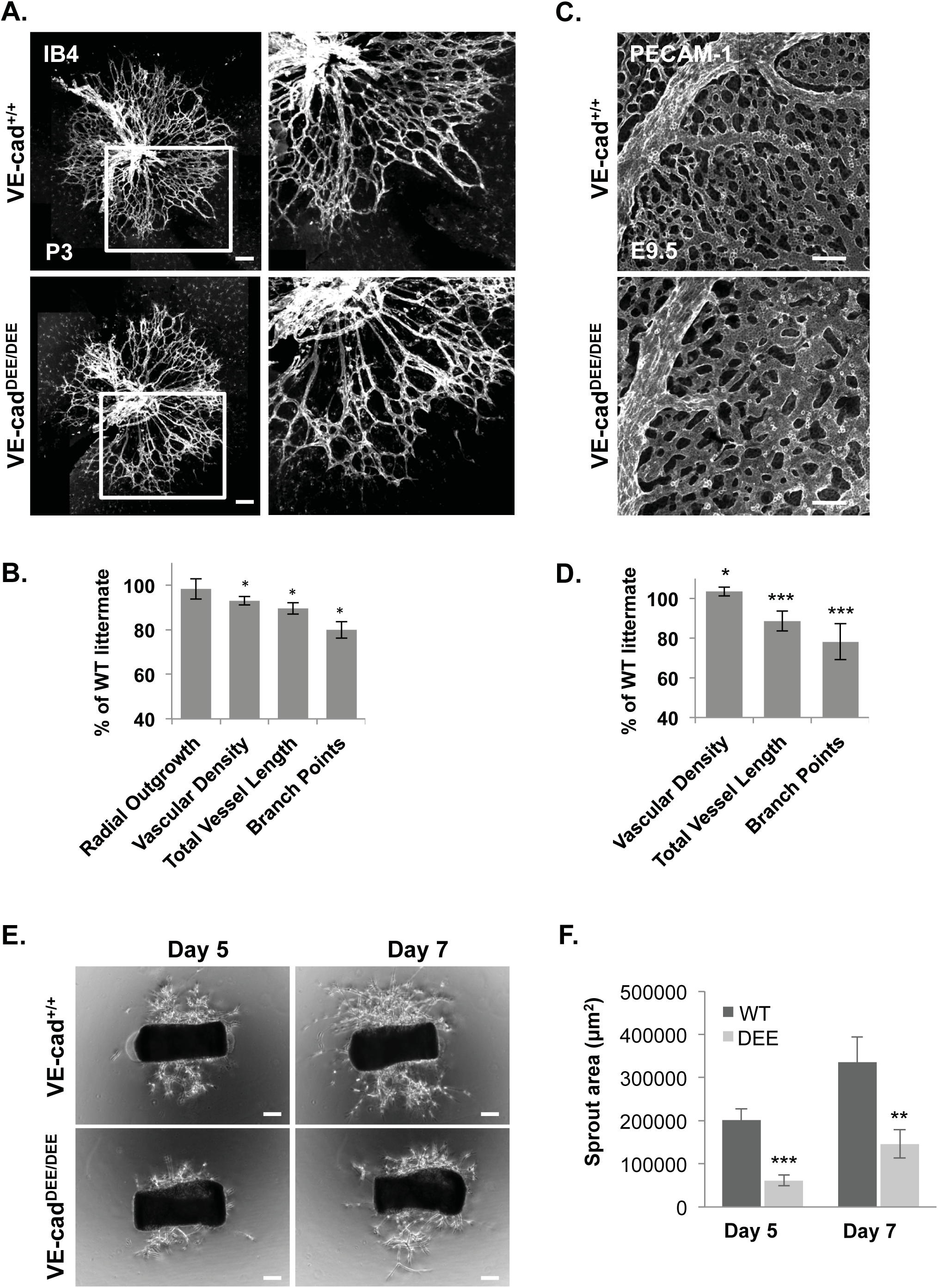
Defects in blood vessel morphology in VE-cad DEE endocytic mutant mice. **A**, Decreased angiogenesis in VE-cad^DEE/DEE^ mutant retinas at P3, as revealed with Isolectin-B4 staining. VE-cad^DEE/DEE^ mutants displayed decreased vessel density and branching at the vessel front compared to VE-cad^+/+^ littermates. Panels on the right show higher magnification of the boxed region in left panels. Scale bar: 100 μm. **B**, Quantitation of the vascular parameters per unit area at the vessel front in P3 VE-cad^DEE/DEE^ retinas. Data are presented as % of WT littermate control and graph represents mean ±SEM. *p<0.05; n=6 independent litters. **C**, Decreased blood vessel formation in VE-cad^DEE/DEE^ mutant yolk sacs at E9.5. Yolk sacs were immunostained for PECAM-1 to visualize blood vessels. Scale bar: 100 μm. **D**, Quantitation of vascular morphology in VE-cad^DEE/DEE^ mutant yolk sacs at E9.5. Data are presented as % of WT littermate control. *p<0.01, ***p<.0001; n=5. **E**, Decreased vessel outgrowth in *ex vivo* aortic ring assays in VE-cad^DEE/DEE^ mutant mice. 1 mm rings from the aortas of adult VE-cad^+/+^ or VE-cad^DEE/DEE^ mice were embedded in Matrigel and analyzed after 5 and 7 days for vessel outgrowth by phase contrast. Scale bar: 150 μm. **F**, Area of vessel outgrowth from aortic rings at the indicated days was quantitated. **p<0.01, ***p<0.001.

To further investigate vascular defects in the VE-cad^DEE/DEE^ mutant, we analyzed blood vessel organization in the developing yolk sac. Following formation of the primitive vascular plexus (E8.5), vessels undergo extensive angiogenic remodeling events that involve nascent vessel sprouting, intussusception, vessel fusion and pruning, leading to a hierarchically ordered network of branched vessels visible by E9.5 (Garcia and Larina, 2014; Udan et al., 2013). Examination of yolk sac whole mount preparations from the VE-cad^DEE/DEE^ mutants revealed normal formation of vitelline and larger diameter vessels, but abnormal microvascular patterning (Figure 6C). In particular, many microvessels were enlarged and appeared to have undergone less angiogenic remodeling than in WT littermates. Quantitation revealed a slight increase in vascular density in the mutants compared to their WT littermates, as well as a significant decrease in the total vessel length and branching (Figure 6D). Thus, vessels are dilated and the network is less complex in the VE-cad^DEE/DEE^ mutants compared to WT littermates. To further test for the presence of sprouting defects in VE-cad^DEE/DEE^ mutants, we performed *ex vivo* aortic ring assays. Rings cut from adult aortas were embedded in Matrigel and the area of sprout outgrowth was quantified after 5 or 7 days of growth. As shown in Figure 6E,F, VE-cad^DEE/DEE^ mutants exhibited significantly reduced network formation at both time points compared with WT littermate controls. Together, these data reveal an essential role for VE-cad endocytosis for endothelial remodeling and sprouting angiogenesis.

### VE-cad endocytosis is required for endothelial polarization and collective cell migration

Sprouting angiogenesis is a form of collective cell movement involving dynamic and continuous interchange between endothelial cells migrating as groups, driven in part by differential adhesion (Bentley et al., 2014). We hypothesized that reduced junction plasticity could inhibit collective migration, which in turn could lead to the observed vessel patterning defects. To test this, we isolated primary endothelial cells from the dermis of early postnatal WT and VE-cad^DEE/DEE^ pups and performed scratch wound migration assays. Analysis of wound closure over a 12-hour period revealed a slower migration rate of VE-cad^DEE/DEE^ endothelial cells compared with WT endothelial cells (Figure 7A,B). We observed no migration defects in VE-cad^GGG/GGG^ mutant endothelial cells, consistent with normal microvessel development observed in these mutant mice (Supplementary Figure 3A,B). Together, these data indicate that VE-cad endocytosis is required for the collective cell movements that occur during sprouting angiogenesis.

**Figure 7.**
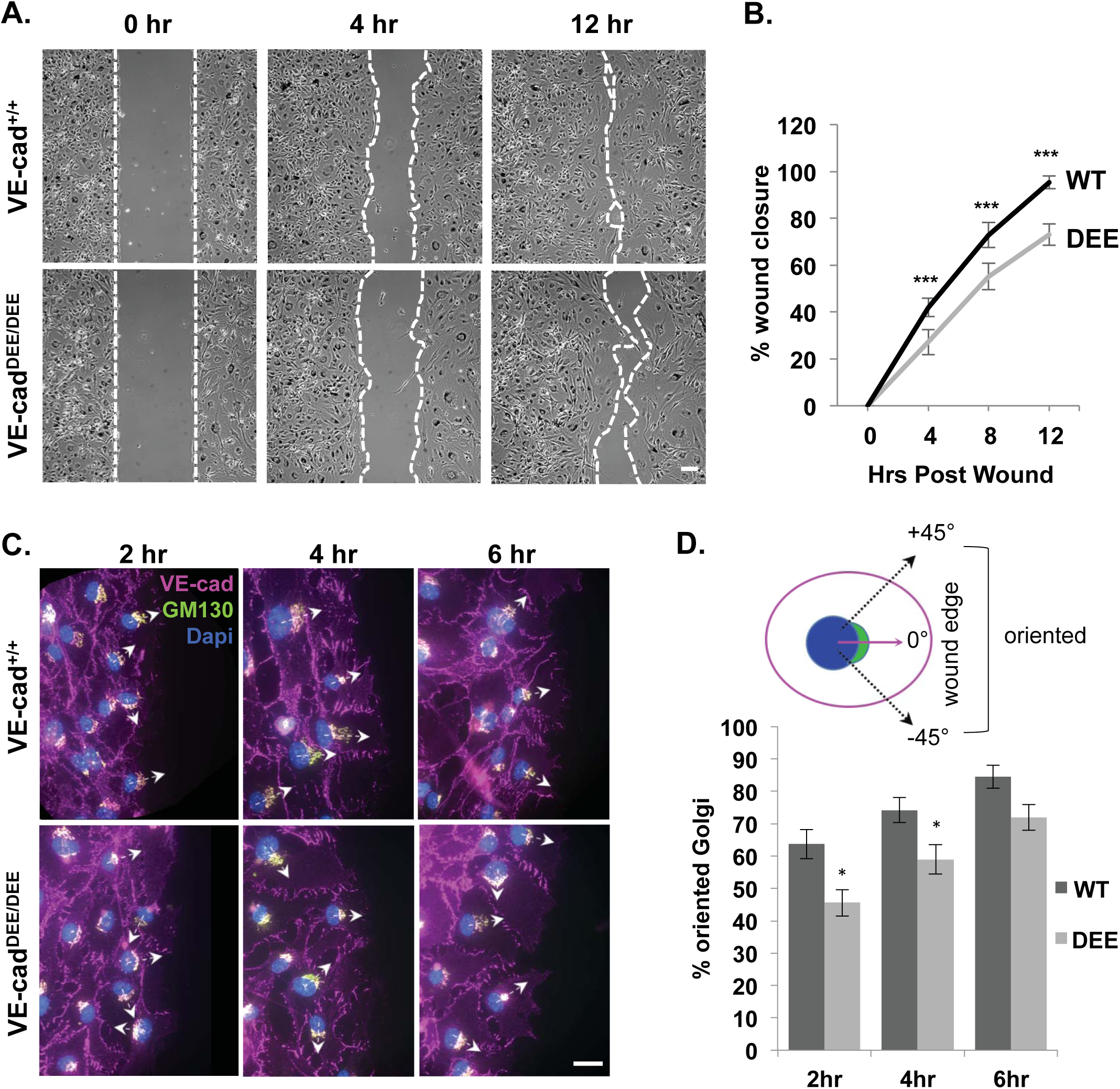
VE-cad DEE endocytic motif is required for endothelial migration and polarization *in vitro*. **A**, Decreased migration of isolated VE-cad^DEE/DEE^ mutant dermal endothelial cells *in vitro*. Scratch-wound assays were performed with primary dermal microvascular endothelial cells isolated from VE-cad^+/+^ and VE-cad^DEE/DEE^ mutant mice. White dashed lines denote scratch borders. **B**, The percentage wound closure by VE-cad^+/+^ and VE-cad^DEE/DEE^ mutant cells was calculated over 12 hours using phase-contrast microscopy. Graph is representative of 4 independent experiments. ***p<0.001. **C**, Decreased polarization in wound edge VE-cad^DEE/DEE^ mutant endothelial cells at the indicated time points post-wounding. Golgi polarity in wound edge cells was analyzed by immunostaining for the Golgi marker GM-130 (green), nuclei (blue) and VE-cad (red). Arrows indicate the nucleus-Golgi polarity axis. **D**, The percentage of cells with their Golgi polarized towards the wound edge in scratch wound assays was determined at 2 hours, 4 hours, and 6 hours post-wounding. A line from the nucleus through the center of the Golgi was drawn and the percentage of cells correctly oriented at ±45° towards the wound edge was calculated. Graph is representative of 4 independent experiments. *p<0.05.

Polarization of the cell motility machinery is a critical first step during collective cell migration. This polarization involves asymmetric membrane trafficking, centrosome/Golgi complex reorientation toward the leading edge, polarized activation of Rho family GTPases, and remodeling of the microtubule and actin cytoskeletons to generate a protrusive front and a retracting rear (Khalil and de Rooij, 2019; Mayor and Etienne-Manneville, 2016). To investigate whether VE-cad endocytosis is required for polarization during collective migration, we examined reorientation of the Golgi apparatus in VE-cad^DEE/DEE^ mutant endothelial cells in response to wounding. Primary WT or VE-cad^DEE/DEE^ mutant endothelial cells were processed for immunofluorescence localization of VE-cad and the Golgi marker, GM-130 together with Dapi at various times after wounding. We defined cells with the Golgi localized within a 90° quadrant between the nuclei and the wound edge as polarized cells (see drawing in Figure 7D). As shown in Figure 7C,D, we observed a slower rate of polarization of the VE-cad^DEE/DEE^ mutant endothelial cells compared to WT cells in response to wounding. After 2 hours, 62% of WT cells were polarized whereas only 45% of DEE mutant cells were polarized. Likewise, 75% of WT cells were polarized after 4 hours, whereas only 52% of mutant cells were polarized. At 6 hours, although the percentage of DEE mutant cells was still slightly decreased compared to WT, this difference was no longer statistically significant. Thus, VE-cad^DEE/DEE^ mutant cells eventually polarize, but at a slower rate than WT cells. We observed similar Golgi reorientation defects in wounded human microvascular endothelial cells expressing the VE-cad DEE→AAA (VE-cadDEE) mutant, but not those expressing wild type VE-cad or the GGG→AAA (VE-cadGGG) mutant (Supplementary Fig 3C,D). These findings suggest that cadherin-mediated polarization does not require p120 binding. These data also suggest that the VE-cadDEE mutant can act dominantly over wild type to suppress polarization. Thus, stabilization of VE-cad on the cell surface inhibits the ability of endothelial cells to polarize in response to directional migration cues.

### VE-cad endocytosis drives actin cytoskeleton remodeling during endothelial polarization

Cadherin-mediated adhesions have previously been shown to regulate cell polarity in wounded monolayers and cell colonies in an actin cytoskeleton-dependent manner (Desai et al., 2009; Dupin et al., 2009). Moreover, remodeling of the actin cytoskeleton is known to play a crucial role in the collective migration of endothelial cells and in the reorganization of endothelial cell junctions (Huveneers and de Rooij, 2013; Vitorino and Meyer, 2008). We hypothesized that expression of the stabilized VE-cadDEE mutant, which interacts with the actin cytoskeleton through C-terminal β-catenin binding (Nanes et al., 2012), may be inhibiting remodeling of the actin network required for polarization. Previous studies have shown that more mature, stable junctions are associated with parallel actin bundles that form a broad band around the cell periphery (Huveneers et al., 2012; Zhang et al., 2005). However, collective cell movements induced by scratch wounding lead to a reduction in junctional actin and the formation of new radial actin bundles (de Rooij et al., 2005; Huveneers and de Rooij, 2013; le Duc et al., 2010; Mangold et al., 2011). Therefore, we examined the organization of the actin cytoskeleton in confluent HUVECs expressing either VE-cadWT or VE-cadDEE. Three hours after wounding, we found that VE-cadWT and VE-cadDEE cells within the monolayer, behind the leader cells at the wound edge, displayed a mix of thick cortical actin bundles and/or thin radial actin bundles (Figure 8A). However, VE-cadDEE mutant cells showed an increased number of cells with thick peripheral actin bundles and a decreased number with radial actin filaments, as compared to VE-cadWT cells (Figure 8A). To quantify this, we applied a threshold mask to the F-actin staining within a field of cells and then measured the intensity and area of the masked regions. We found that although VE-cadDEE expressing cells displayed no difference in the overall fluorescence intensity of F-actin, they did have less F-actin positive area than VE-cadWT cells (Figure 8B,C). This suggests that although the total levels of F-actin are unchanged in VE-cadDEE mutant cells, VE-cadDEE expression inhibits the ability of the cell to remodel actin in response to wounding.

**Figure 8.**
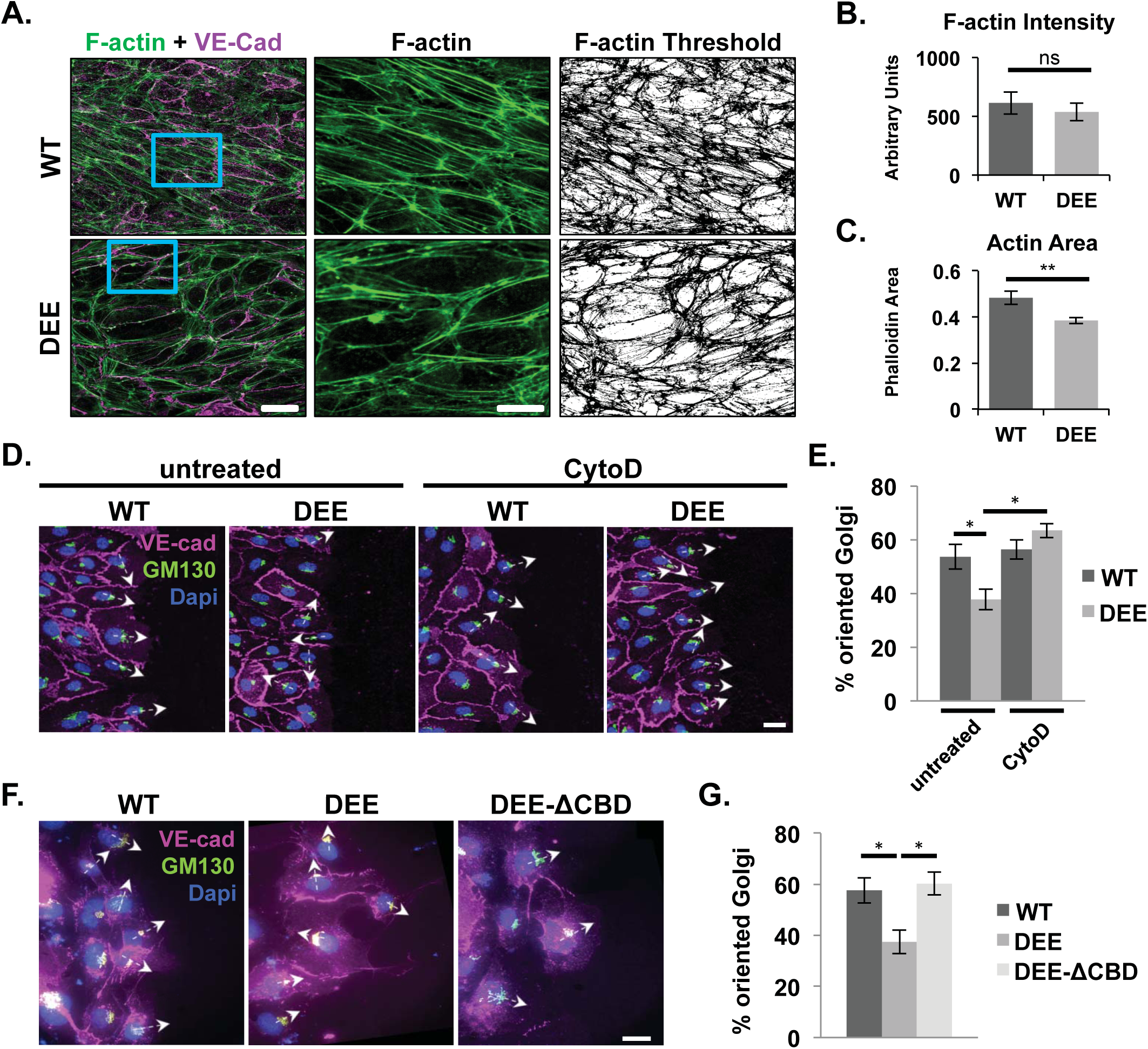
Endothelial polarization driven by VE-cad endocytosis requires actin remodeling. **A**, Rescue of Golgi reorientation defects in VE-cadDEE expressing cells after Cytochalasin D treatment and washout. Cells were transduced with RFP-tagged VE-cadWT or VE-cadDEE adenovirus. Immediately following wounding, cells were either incubated for 2 hours with 1 μm Cytochalasin D followed by 3 hour washout, or left untreated for 3 hours following wounding. Cells were then fixed and immunostained for RFP (red), GM-130 (green) and nuclei (blue) to visualize reorientation of the Golgi towards the wound edge. Arrows indicate the nucleus-Golgi polarity axis. **B**, Graph shows the average percentage ± SEM of RFP-positive cells with a polarized Golgi apparatus (in a 90° quadrant towards the wound edge) 3 hours after washout (treated) or wounding (untreated). Graph represents average of 4 independent experiments. *p<0.05. **C**, Rescue of polarization defects in VE-cadDEE expressing cells by deletion of the catenin-binding domain. HUVEC endothelial cells were transduced with lentivirus coding for the indicated RFP-tagged VE-cad proteins. Cells were fixed 2 hours after wounding and immunostained for GM-130 (green) and nuclei (blue) to visualize reorientation of the Golgi towards the wound edge. The RFP signal is shown in red. Arrows indicate the nucleus-Golgi polarity axis. **D**, Graph shows the average percentage of RFP-positive cells with their Golgi apparatus polarized in a 90° quadrant towards the wound edge 2 hours after wounding. Graph represents average ±SEM calculated from 3 independent experiments. *p<0.05.

To test this possibility, we attempted to rescue the VE-cadDEE-induced polarization defect by disrupting the actin cytoskeleton with Cytochalasin D for two hours, followed by a three hour washout. This treatment should release the inhibition of actin remodeling due to VE-cadDEE expression. Indeed, we observed normal Golgi polarization in the VE-cadDEE mutant cells following Cytochalasin D treatment and washout (Figure 8D,E). These data suggest that VE-cad endocytosis controls polarization by promoting actin-dependent remodeling during collective migration.

To confirm that VE-cad endocytosis-dependent polarization requires actin remodeling, we next generated a VE-cadΔCBD-DEE compound mutant. This mutant contains both the DEE mutation, as well as a deletion of the C-terminal catenin binding domain (CBD) domain, which links VE-cad to the actin cytoskeleton. Thus, while the VE-cadΔCBD-DEE mutant forms stable adhesions, they are largely uncoupled from the actin cytoskeleton. We then analyzed Golgi reorientation in VE-cadΔCBD-DEE expressing cells. Interestingly, cells expressing the VE-cadΔCBD-DEE compound mutant exhibited normal polarization after wounding (Figure 8F,G). Two hours after wounding, 60.3±4.4% of VE-cadΔCBD-DEE expressing cells were polarized whereas only 37.5±4.9 % of VE-cadDEE mutant cells were polarized. Together, these data suggest that cadherin endocytosis is required for actin remodeling that promotes polarization during collective migration. Furthermore, these findings indicate that the functions of the p120 and β-catenin binding domains of the cadherin tail are integrated to modulate cell polarity and directional movement.

## Discussion

The findings presented here advance two important aspects of classical cadherin biology. First, although p120 participates in numerous cellular activities, we use multiple experimental approaches to demonstrate that the primary function of p120 is to stabilize cell surface cadherin. Indeed, mice harboring a VE-cad mutant (VE-cad^ΔJMD/ΔJMD^) unable to bind p120 but lacking the DEE endocytic motif are apparently normal even though endothelial cells in these animals lack junctional p120. Similarly, the lethality of the endothelial p120 null phenotype can be largely rescued by expressing a VE-cad mutant deficient in endocytic activity (VE-cad^DEE/DEE^). Thus, p120 can be rendered dispensable in the context of a stabilized cadherin. Secondly, we utilized the VE-cad^DEE/DEE^ endocytic mutant to reveal that cadherin endocytosis is required for normal tissue morphogenesis. Surprisingly, VE-cad endocytosis regulates endothelial migration by permitting cell polarization at the onset of collective cell migration. Thus, the cadherin-p120 complex dictates the plasticity of endothelial junctions to govern key functions of vascular endothelial cells, including endothelial cell migration, microvascular patterning, and the acquisition of the vascular barrier.

Previous studies in Drosophila indicate that p120 is not required for survival (Myster et al., 2003; Pacquelet et al., 2003). Subsequent to these initial studies in flies, a series of p120 conditional null mouse mutants confirmed that p120 is essential for cadherin function in most vertebrate tissues, including vascular endothelial cells (Cadwell et al., 2016; Oas et al., 2010). However, these previous vertebrate studies do not determine if p120 is required at the cadherin complex or if p120 is performing other essential functions independent of cadherin binding. Our analysis of the VE-cad^GGG/GGG^ mutant in Figure 1 demonstrates that p120 binding to the cadherin tail is critical for cadherin stability *in vivo* in a vertebrate model system. Although p120 was still present at normal levels in VE-cad^GGG/GGG^ mutant endothelial cells, it failed to localize to cell junctions. VE-cad levels were reduced in these animals and vessel stability was compromised as demonstrated by the presence of embryonic and postnatal hemorrhaging. In addition, these animals were highly susceptible to challenges to the vascular barrier and displayed large increases in lung permeability in response to LPS treatment. Thus, in this vertebrate model system, p120 binding to cadherin is required for normal cadherin expression levels, adherens junction integrity, and overall viability.

Our previous studies demonstrated that the core p120 binding domain of VE-cad harbors a three amino acid endocytic signal (DEE) (Nanes et al., 2012). We hypothesized that cadherin JMD deletions that eliminated both p120 binding and the DEE endocytic motif would result in a cadherin that is deficient in p120 binding but simultaneously stable at the cell surface. Using the CRISPR-Cas system, we generated animals with an 11 amino acid deletion that completely eliminated p120-binding and the DEE endocytic motif (VE-cad^ΔJMD/ΔJMD^). In Figure 3, analysis of junctional p120 in aortic endothelial cells of this mutant revealed that p120 was completely absent from junctions while VE-cad levels were identical to wild type controls. Remarkably, these animals were viable and lacked the hemorrhaging and runting phenotypes associated with the VE-cad^GGG/GGG^ mutation. These results indicate that a classical cadherin can be rendered p120-independent by cadherin mutations that eliminate endocytic activity.

To further examine the requirement for p120, we tested whether a VE-cad endocytic mutant (VE-cad^DEE/DEE^) could rescue the endothelial p120 null phenotype. Indeed, most p120 null animals expressing the VE-cad^DEE/DEE^ mutation survived, in contrast to animals expressing WT VE-cad, which exhibited almost complete embryonic lethality (Figure 5). Furthermore, VE-cad levels at cell-cell borders in VE-cad^DEE/DEE^ mutants lacking p120 were normal, confirming that the VE-cad^DEE/DEE^ mutant is stable in the absence of p120 binding. However, it should be noted that often the surviving Tie2-Cre; VE-cad^DEE/DEE^; p120^flox/flox^ mice were smaller than their Cre negative littermates, and occasionally died within a few weeks after weaning. There are at least two possible explanations for this outcome. One likely possibility is that p120 is also required for N-cad regulation. N-cad levels are reduced in endothelial cells lacking p120 (Davis et al., 2003; Ferreri et al., 2008; Oas et al., 2010), and several studies indicate that N-cad is essential for early vascular development and pericyte recruitment (Luo and Radice, 2005; Tillet et al., 2005). Second, p120 is known to perform other, cadherin-independent functions such as the regulation of Rho GTPases and Kaiso mediated transcriptional regulation (Dunach et al., 2017; Kourtidis et al., 2013). Thus, the complete loss of p120 is likely to be more severe than the VE-cad^GGG/GGG^ mutation because multiple cadherins are affected by loss of p120 and because p120 carries out cadherin independent roles.

Several studies have reported that abrogation of VE-cad expression leads to vessel hyper-sprouting in mice (Gaengel et al., 2012) and zebrafish (Abraham et al., 2009; Montero-Balaguer et al., 2009). However, we did not observe an aberrant patterning or sprouting phenotype in VE-cad^GGG/GGG^ mutants. Although blood spots were increased by five-fold in the VE-cad^GGG/GGG^ mutant retina, blood vessel density and patterning were similar to wild type. One possibility is that endothelial cells need a threshold level of VE-cad to prevent excess sprouting. This idea is supported by a VE-cad knockdown study in zebrafish (Montero-Balaguer et al., 2009). Although high concentrations of VE-cad morpholinos prevented the establishment of reciprocal contacts between vessels, leading to increased sprouting, low doses of VE-cad morpholinos led to vascular fragility, hemorrhages and increased permeability, a phenotype reminiscent of VE-cad^GGG/GGG^ mutants. Therefore, the mechanisms by which VE-cad controls sprouting behavior are likely complex and dependent upon the levels of VE-cad present.

It is interesting to note that the severity of the vascular integrity phenotype in various VE-cad mutants directly correlates with the levels of VE-cad present at cell junctions. In homozygous null mice, a complete lack of VE-cad leads to severe vascular defects and early embryonic death due to the regression and disintegration of nascent blood vessels (Carmeliet et al., 1999; Crosby et al., 2005; Gory-Faure et al., 1999). However, heterozygous null mice, which display a 50% reduction in VE-cad levels, were reported to have no obvious defects (Carmeliet et al., 1999; Gory-Faure et al., 1999). The VE-cad^GGG/GGG^ mutants reported here display a 70% reduction in VE-cad levels and have a moderately severe phenotype (Figure 1). The VE-cad^ΔJMD/ΔJMD^ and VE-cad^DEE/DEE^ mutants have normal VE-cad levels and no hemorrhaging or lethality. We would therefore predict that any further reduction of functional VE-cad in VE-cad^GGG/GGG^ mutants would lead to a more severe phenotype. Consistent with this possibility, we found that VE-cad^GGG/STOP^ mutants all die embryonically. Together, these data suggest that a threshold level of VE-cad between 30-50% normal levels is required for normal vascular integrity and survival.

It is also interesting to note that the VE-cad^GGG/GGG^ mutants that survived the first few postnatal days usually survived into adulthood. We did not observe an increase in the death rate of older (P7 or later) VE-cad^GGG/GGG^ mutants, despite having reduced VE-cad levels. Furthermore, the reduced body weight of VE-cad^GGG/GGG^ mutants was more pronounced at 3 weeks as compared to 6 weeks (Figure 1). Together, this suggests that the maintenance of normal VE-cad levels is most critical during the early stages of blood vessel morphogenesis, and that other mechanisms likely compensate for reduced VE-cad levels during later developmental stages. It is unlikely that N-cad was compensating for the loss of VE-cad in VE-cad^GGG/GGG^ mutants because we failed to observe N-cad at cell junctions in the mutant aortas. In addition, β-catenin levels at cell junctions paralleled the reduction of VE-cad at junctions. Thus, once the adherens junctions are formed, they can likely be maintained despite loss of the cadherin. This finding is supported by work from Frye et al. showing that induced deletion of the VE-cad gene in adult (7 week old) mice led to increased vascular permeability in the heart and lungs but no obvious vascular abnormalities or lethality (Frye et al., 2015).

A number of studies have implicated cadherin endocytosis in the regulation of tissue morphogenesis in both flies and mice. However, a limitation of previous studies is the reliance on broad inhibition of endocytic or recycling pathways to modulate cadherin trafficking (Cadwell et al., 2016; Ratheesh and Yap, 2010). The generation of the VE-cad^DEE/DEE^ mouse line allowed us to directly test the role of cadherin endocytosis in vertebrate morphogenesis. The formation of large vessels was grossly normal in these mutants. However, as shown in Figure 6, VE-cad^DEE/DEE^ mutants displayed defects in vessel density, length, and branching during postnatal retina angiogenesis. These mutants also exhibited decreased vessel length and branching in the yolk sac microvasculature, as well as less neovessel outgrowth in *ex vivo* aortic rings. Endothelial cells isolated from the VE-cad^DEE/DEE^ mutant mice were defective in collective migration as assessed using scratch wound assays (Figure 7). Similar collective cell migration defects were observed in microvascular endothelial cells expressing the VE-cadDEE mutant but not in cells expressing WT VE-cad or the VE-cadGGG mutant. It is interesting to note that we observed no increase in VE-cad levels in the VE-cad^DEE/DEE^ mutant, either *in vitro* or *in vivo*. This finding suggests that cells utilize unknown mechanisms to limit total cadherin cell surface levels when turnover rates are experimentally reduced. Furthermore, our results suggest that VE-cad endocytic rates rather than overall cadherin expression levels are the critical determinant controlling collective migration.

Analysis of migratory activity of cells expressing the VE-cad^DEE/DEE^ mutant revealed that VE-cad endocytosis is required for endothelial cell polarization at the onset of collective cell migration. This defect was observed both in cells isolated from VE-cad^DEE/DEE^ mutant mice and in endothelial cells exogenously expressing the VE-cadDEE mutant. Importantly, deletion of the β-catenin binding domain from the tail of the VE-cadDEE mutant relieved these polarity and migration defects caused by the endocytic mutation, as shown in Figure 8. Furthermore, disruption of the actin cytoskeleton using Cytochalasin D followed by washout, rescued the polarity defects in cells expressing the VE-cadDEE mutant. Together, these data indicate that linkage to the actin cytoskeleton is required for the inhibitory activity of the DEE mutant. We also observed altered organization of the actin cytoskeleton in VE-cadDEE expressing cells. These cells exhibited an increase in thick, parallel bundles around the cell periphery and decreased radial actin fibers, a pattern characteristic of stable junctions. Classical cadherins have been implicated in polarization and collective cell movements by directing migration machinery and protrusive activity away from cell-cell contacts and towards the leading edge (Desai et al., 2009; Dupin et al., 2009; Theveneau et al., 2010; Weber et al., 2012). Although the mechanisms that promote cadherin-mediated polarization are poorly understood, they may involve anisotropic distribution of cadherin molecules or asymmetric signaling from cadherin-mediated junctions, which could be disrupted in the stabilized VE-cad^DEE/DEE^ mutant (Dorland et al., 2016; Hayer et al., 2016; Mayor and Etienne-Manneville, 2016). Previous studies have implicated cadherin treadmilling along lateral borders, as well as endocytosis and recycling, in the process of collective cell migration (Peglion et al., 2014). Also, Cao et al. found that VEGF-induced polarized cell elongation during endothelial collective migration involves a reduction in the relative concentration of VE-cad at junctions. This reduction in cadherin triggers the formation of small, actin-driven junction-associated intermittent lamellipodia (JAIL), which are associated with increased migration (Abu Taha et al., 2014; Cao et al., 2017). Combined with the results presented here showing VE-cad endocytosis drives actin dynamics during collective migration, it is attractive to speculate that VE-cad endocytosis may promote JAIL formation that, in turn, is required for polarization and collective migration. Additional studies will be required to fully understand how cadherin dynamics are modulated by endocytic pathways to permit polarized endothelial cell migration in normal development and pathological circumstances such as tumor angiogenesis.

## Materials and Methods

### Mice

Tie2-Cre mice (#004128) and VE-cad-Cre mice (#017968) were obtained from Jackson Laboratory and have been described previously (Chen et al., 2009; Koni et al., 2001). Mice with LoxP sites in introns 2 and 8 of the p120 gene were described previously (Davis and Reynolds, 2006). We generated point mutant mice with alanine substitutions in either DEE resides (amino acids 646-648) or GGG residues (amino acids 649-651) using CRISPR/Cas9 genome editing in conjunction with the Mouse Transgenic and Gene Targeting Core Facility at Emory. The CRISPR guide RNA (gRNA) sequence CAGTTGGTCACTTACGATG used for both mutants contains at least 3 base pair mismatches against any other targets in the mouse genome. Long, single stranded oligonucleotides containing the indicated point mutations at either the DEE or GGG site were injected into the cytoplasm of one-cell zygotes together with Cas9 mRNA and the gRNA. Founder pups with DEE or GGG point mutations generated by homology-directed repair with the donor oligo were identified by PCR/restriction fragment length polymorphism (RFLP) analysis of genomic DNA. Primers CM13, 5’-CTGGTCCCATGAACCTGTCT-3’ and CM14, 5’-GCGCACAGAATTAAGCACTG-3’, were used to amplify a 212bp product, which was digested with Fnu4H1. Both DEE and GGG mutant alleles contain a new Fnu4H1 site absent in the wild type allele PCR product. The correct mutations were also confirmed by Sanger sequencing of genomic DNA. The ΔJMD mouse strain containing a deletion of amino acid residues 642-653 (LVTYDEEGGGE) was generated as a result of non-homologous end joining (NHEJ) during the process of making the DEE point mutant strain. The STOP allele of VE-cad contains a TGA stop codon at amino acid 647, and an aspartic acid to leucine point mutation at amino acid 646, and was also generated by non-homologous end joining (NHEJ) while making the DEE point mutant strain. All procedures were performed in accordance with NIH guidelines and the US Public Health Service’s Guide for the Care and Use of Laboratory Animals and were approved by the IACUC of Emory University, which is accredited by the American Association for Accreditation of Laboratory Care (AAALC).

### Aorta *en face* immunostaining

Six-to eight-week-old mice were euthanized by CO_2_ inhalation and immediately perfused through the left ventricle with Dulbecco’s phosphate-buffered saline (DPBS) containing 10,000 U/L heparin followed by fresh 4% paraformaldehyde in DPBS for 8 minutes. The thoracic aorta was then carefully removed and cleaned of fat in a Petri dish containing DPBS. The aorta was then opened to expose the lumen and cut into pieces for immunostaining. Aortas were incubated two times in permeabililzation buffer (0.25% Triton X-100 in DPBS) for 20 minutes at room temperature followed by incubation in blocking buffer (10% normal goat serum plus 0.25% Triton X-100 in DPBS) for 2 hours at room temperature. Samples were then incubated overnight at 4°C with primary antibodies diluted in blocking buffer. Primary antibodies were: rat anti-VE-cad (BV13, eBioscience **#**14144181, 1:500); rabbit anti-p120 (S-19, Santa Cruz Biotechnology, Inc #sc-1101, 1:250); mouse anti-beta-catenin (BD Biosciences #610153, 1:250); mouse anti-N-cadherin (BD Biosciences #610920, 1:250). Aortas were then washed four times in DPBS for 15 minutes each and subsequently incubated with secondary antibodies diluted in blocking buffer for two hours at room temperature in the dark. Secondary antibodies were: Alexa Fluor 555 goat anti-rat IgG, (Invitrogen #A-21434, 1:3000); Alexa Fluor 488 goat anti-rabbit IgG (Invitrogen #A-11008, 1:3000); Alexa Fluor 488 goat anti-mouse IgG (Invitrogen #A-11029, 1:3000). Samples were washed four times in DPBS and then mounted on glass slides with intima side up using ProLong Gold (Invitrogen #P36930). Aortas were imaged with either a Zeiss LSM510 Meta confocal microscope with ZEN 2009 software or an Olympus BX61WI upright confocal microscope with Olympus Fluoview v4.2 acquisition software. At least 5-6 fields of view were imaged from each mutant and wild type littermate control. At least four separate litters were analyzed for each mutant, and all comparisons were made in similar regions of the aorta between mutant and wild type. Images were analyzed using Nikon NIS-Elements AR version 4.40 software and processed using ImageJ Fiji. The protein expression levels at cell-cell borders were quantitated by creating a mask of the borders in each field based on anti-VE-cad immunostaining, and measuring the average fluorescent intensity of the protein of interest within this mask. Protein levels in wild type were set to 100% for each litter and the average % decrease in the indicated homozygous mutant littermate was quantitated. Statistics were computed using GraphPad Prism 7 and the Mann-Whitney *U* test used to evaluate significance.

### Retina angiogenesis

P3 mice were euthanized by decapitation. For lectin staining, eyes were enucleated and immediately fixed overnight at 4°C in 4% paraformaldehyde in DPBS. Fixed eye cups were photographed for blood spots on a Nikon SMZ745T stereo Microscope with a Nikon DS-Fi3 Digital Camera + DS-L4 control unit. Blood spot area was quantitated using ImageJ Fiji software, and a *t* test was used to evaluate significance. Eyes were then washed five times in DPBS and retinas were dissected out and flattened by making four radial incisions. Retinas were then incubated in blocking buffer (0.3% Triton X-100 plus 10 mg/mL BSA in DPBS) overnight at 4°C. The next day, retinas were incubated overnight at 4° C with fluorescein labeled Griffonia Simplicifolia Lectin I isolectin B4 (GSL I-B4; Vector Laboratories; Burlingame, CA #FL-1201) diluted 1:50 in blocking buffer. Samples were then washed and mounted on glass slides using ProLong Gold (Invitrogen #P36930). Images were acquired on a Zeiss LSM510 Meta confocal microscope and stitched together using the stitching plugin from ImageJ Fiji. Vascular density, total vessel length and branchpoints per unit area at the vascular front were quantitated using the Angiotool software (National Institutes of Health National Cancer Institute, Gaithersburg, MD) (Zudaire et al., 2011). Radial outgrowth was analyzed by measuring the radial distance from the optic nerve head to the vascular front at the retinal periphery. A paired *t* test between wild type and mutant littermates was used to evaluate significance.

### Embryo and yolk sac analysis

For timed pregnancies, the morning of the plug was designated as E0.5 and the day of birth postnatal day 0 (P0). Whole unfixed embryos were harvested at E12.5 and examined and photographed on a Nikon SMZ745T stereo Microscope with a Nikon DS-Fi3 Digital Camera + DS-L4 control unit. For yolk sac analysis, wild type and mutant littermates were harvested at E9.5 or E12.5 and yolk sacs were removed and fixed in 4% paraformaldehyde in DPBS for 1.5 hours at room temperature. This was followed by two DPBS washes. Samples were then incubated in blocking buffer (10% normal goat serum plus 0.1% Triton-X-100 in DPBS) for 2 hours at room temperature followed by overnight incubation at 4°C with anti-PECAM-1 (MEC13.3, BD Bioscience #550274, 1:250) diluted in blocking buffer. The following day, four washes in DPBS plus 0.1% Triton X-100 were performed, followed by incubation with Alexa Fluor 555 goat anti-rat IgG secondary antibody (Invitrogen #A-21434, 1:3000) for two hours at room temperature in the dark. Following four more washes, samples were mounted using ProLong Gold (Invitrogen #P36930). Overlapping images of the entire E9.5 yolk sacs were taken either on a Zeiss LSM510 Meta confocal microscope or an Olympus BX61WI upright confocal microscope. Images were stitched together using the stitching plugin from ImageJ Fiji. Vascular density, total vessel length and branchpoints at comparable regions of the stitched yolk sacs were analyzed using Angiotool software. Quantitation was based on 4-7 pairs of images within the yolk sac for each set of embryos (paired for corresponding locations within the yolk sac), and a paired *t* test between location-paired images from the wild type and mutant was used to evaluate significance.

### p120 conditional knockout rescue

To analyze rescue of VE-cad levels in p120CKO mice by the DEE mutant allele, matings were set up between VE-cad-Cre^+^; VE-cad^DEE/+^; p120^flox/flox^ and VE-cad^DEE/+^; p120^flox/flox^ mice to generate both VE-cad-Cre^+^; VE-cad^+/+^; p120^flox/flox^ and VE-cad-Cre^+^; VE-cad^DEE/DEE^; p120^flox/flox^ mice for analysis. VE-cad-Cre mediated deletion of p120 resulted in mosaic deletion of p120, leading to fields of view with both p120-positive (p120^+^) and p120-negative (p120^−^) cells. To quantitate VE-cad levels, cell-cell borders between two p120^+^ or two p120^−^ cells were traced in ImageJ Fiji and the average VE-cad levels were measured. The p120^+^ and p120^−^ borders were from the same field and the average VE-cad level at the p120^+^ cell-cell borders in each sample was normalized to 100%. At least 5-10 fields of view (at least ten 120^+^ and ten p120^−^ borders per field) from three independent experiments were analyzed for each genotype. A *t* test was to evaluate significance. To analyze lethality rescue by the VE-cad DEE mutant allele, matings were set up between male Tie2-Cre; VE-cad^DEE/+^; p120^flox/flox^ mice and female VE-cad^DEE/+;^ p120^flox/flox^ mice and surviving offspring were genotyped between P6 and P8.

### Aortic ring assay

Aortic rings were performed as described (Baker et al., 2011). Briefly, mice were sacrificed by CO_2_ inhalation and the thoracic aorta was dissected out under sterile conditions and transferred to a Petri dish containing cold Opti-MEM I Reduced-serum Medium + GlutaMAX-I (Gibco, #51985-026). Extraneous fat was removed and blood was gently flushed from the vessel using a 27-G needle fixed to a 1-mL syringe. Using a scalpel, ~1 mm wide aortic rings were cut and embedded in matrigel (BD Biosciences, #354230) in a 24 well plate. Aortic rings were incubated for 5-9 days at 37°C and 5% CO_2_ in the above Opti-MEM medium supplemented with 100 U/mL penicillin and 100 μg/mL streptomycin (Gibco, #15140-122) and 2.5% (v/v) fetal bovine serum. Media was removed every other day and replaced with 1 mL of fresh media. Sprout outgrowth was imaged by phase microscopy using a Biotek Lionheart FX microscope equipped with a Sony ICX285 CCD camera and a 4X objective. The total area of vessel outgrowth minus the area of the ring was quantitated by a blinded observer using ImageJ Fiji software and results are representative of four independent experiments from separate litters, with at least 8-10 rings analyzed per experiment per animal. A *t* test was used to evaluate significance.

### *In Vivo* lung permeability assays

Lung permeability was measured using Evans Blue dye leakage (Radu and Chernoff, 2013). Briefly, 2-3 month old mice received intraperitoneal injections of either normal phosphate-buffered saline (DPBS) or 18 mg/kg of body weight *E. coli* Lipopolysaccharides (LPS) (Sigma #L4391, lot#043M4089V) in DPBS. After six hours, mice were injected intraorbitally with 100 μl of 1% Evans Blue solution (MP Biomedicals lot# QR12404) in DPBS. Evans Blue was allowed to circulate for 15 minutes and mice were transcardially perfused with 50 mL of DPBS. The lungs were then removed and Evans blue was extracted by incubation with formamide at 55°C overnight. Dye concentration was quantitated spectrophotometrically in the supernatant at 620 nm and normalized to the dry weight of the lung. Statistics were computed using GraphPad Prism 7 and significance was evaluated using two-way ANOVA.

### Cell Culture

Primary mouse dermal endothelial cells were obtained using methods previously described (Oas et al., 2010). Briefly, skins from early postnatal pups (P5-P7) were isolated and enzymatically dissociated using 2 mg/mL collagenase type I (Worthington Biochemical Corp, Lakewood, NJ #LS004196) followed by trituration with a cannula. Endothelial cells were then purified by a 10-minute incubation in suspension with magnetic sheep anti-rat Dynabeads (Invitrogen, Carlsbad, CA) coated with rat anti-mouse PECAM-1 (rat; clone MEC 13.3 BD Biosciences #553369). Magnetically sorted cells were then plated in flasks coated with 0.1% gelatin and grown to confluency. A second round of purification was then performed with anti-rat Dynabeads coated with rat anti-mouse ICAM-2 (rat clone 3C4; BD Biosciences #553325). Cells were cultured in Endothelial Cell Growth Medium MV2 (PromoCell, Heidelberg, Germany #C-22022). The endothelial identity of the cells was confirmed by immunofluorescence microscopy with antibodies to endothelial markers PECAM-1 and VE-cadherin. Greater than 95% purity was routinely observed in the preparations. Freshly isolated human umbilical vein endothelial cells (HUVEC) were cultured in Endothelial Cell Growth Medium (PromoCell, Heidelberg, Germany #C-22010). Primary human dermal microvascular endothelial cells (MEC) were isolated from neonatal foreskin and cultured on 0.1% gelatin-coated culture dishes in 0.1% gelatin-coated culture dishes in EGM-2 MV media (Lonza #cc-3202). Both HUVEC and MEC were used before passage five in all experiments and the endothelial identity of the cells was confirmed by immunofluorescence microscopy with antibodies to endothelial markers PECAM-1 and VE-cadherin.

### Western Blotting

Isolated dermal endothelial cells were lysed in Laemmli sample buffer and proteins were separated on a 7.5% Mini-Protean TGX precast gel (#456-1025, Bio-rad) using Tris/glycine/SDS running buffer (#161-0732, Bio-rad) and transferred to a nitrocellulose membrane (#88018, Thermo Scientific). Western blots were developed with chemiluminescence HRP substrate (#RPN2106 GE Healthcare). Antibodies used are goat anti-VE-cad (C-19; Santa Cruz Biotechnology, Inc #sc-6458; 1:500), rabbit anti-p120 (S-19; Santa Cruz Biotechnology, Inc #sc-1101; 1:750), mouse anti-beta-catenin (BD Biosciences #610153; 1:1,000), mouse anti-N-cadherin (BD Biosciences #610920; 1:1,000) and rabbit beta-actin (clone D6A8; Cell Signaling Technology #8457S; 1:1,500). The chemiluminescent blots were imaged with the ChemiDoc MP imager (Bio-Rad). Densitometric analysis was performed using ImageJ Fiji software.

### Wounding assay

Primary mouse endothelial cells were plated in growth medium in Ibidi 35 mm Culture-Insert 3-well wounding assay dishes (Ibidi, Madison, WI #80366) and grown to confluency. Cells were then starved overnight in EBM^TM^-2 basal medium (Lonza cc-3156) containing 1% FBS. The following morning, the silicone insert was removed leaving a 500 μm wound. Each field was imaged by phase contrast with a 10X objective at initiation of wound closure and the indicated time points to monitor wound closure. Images were obtained using a Nikon Eclipse Ti-E Inverted Microscope equipped with a motorized stage and a Hamamatsu C11440-22CU Digital Camera and NIS-Elements software version AR4.40.00. Wound area was measured using ImageJ Fiji software and a *t* test was used to evaluate significance.

### Golgi reorientation

Wounding assays with primary mouse cells were performed as described above. Cells were fixed at the indicated time point with 4% paraformaldehyde in DPBS for 10 minutes at room temperature. Cells were then washed with DPBS and permeabilized in 0.1% Triton X-100 in phosphate-buffered saline for 10 minutes. Cells were then incubated in blocking buffer (0.1% Triton X-100 plus 10% normal goat serum in DPBS) for 20 minutes followed by overnight incubation in primary antibodies at 4°C (mouse anti-GM-130, BD Transduction laboratories #610822 1:250; rat anti-VE-cad BV13, ThermoFisher Scientific **#**14-1441-82 1:500) diluted in blocking buffer. Cells were then incubated with an Alexa Fluor 488 goat anti-mouse IgG antibody (Invitrogen #A-11029, 1:3000) together with Alexa Fluor 555 goat anti-rat IgG (Invitrogen #A-21434, 1:3000) for 1 hour at room temperature and mounted using ProLong Gold with Dapi (Invitrogen; #P36932). HUVEC were plated on gelatin-coated Ibidi wounding assay dishes (described above) or plated on gelatin-coated coverslips in 4-well dishes (Thermo Scientific Nunc, #144444) and infected with the indicated RFP-tagged lentivirus or adenovirus constructs. Three days after infection (for lentivirus) or 8 hours after infection (for adenovirus), cells were starved overnight in EBM^TM^-2 basal medium (Lonza cc-3156) containing 1% FBS. The following morning, inserts were removed (or cells were scratched with a pipette tip) and cells were allowed to polarize for 2-3 hours as indicated. Cells were then fixed, immunostained for GM-130 and RFP (rabbit polyclonal anti-RFP; Rockland #600-401-379) and imaged. For Cytochalasin D experiments, HUVEC infected with VE-cadWT-RFP or VE-cadDEE-RFP adenovirus were treated for 2 hours with 1μM Cytochalasin D (Sigma #C8273) immediately after wounding, washed 3 times in DPBS, and allowed to polarize in complete medium for 3 hours following washout. Untreated VE-cadWT-RFP and VE-cadDEE-RFP cells were allowed to polarize for 3 hours following wounding. Human MEC plated on gelatin-coated coverslips in 4-well dishes were infected in the morning with the indicated RFP-tagged adenovirus constructs and starved overnight. The following morning (24 hours after infection), cells were wounded with a pipette tip and allowed to polarize for 6 hours. Cells were then fixed, immunostained for GM-130 and RFP and imaged. Mouse cell and HUVEC images were captured using a Nikon Eclipse Ti-E Inverted Microscope equipped with a motorized stage, a 60x/1.49NA oil immersion lens and a Hamamatsu C11440-22CU Digital Camera using NIS-Elements software version AR4.40.00. MEC microscopy was performed using an epifluorescence microscope (DMRXA2, Leica) equipped with 63x/1.32NA oil immersion objective, narrow band pass filters, and a digital camera (ORCA-ER C4742–80, Hamamatsu Photonics). MEC images were captured using Simple PCI software (Hamamatsu Photonics). Wound edge cells with the Golgi apparatus localized within the 90° angle in front of the nucleus, facing the wound axis, were quantified as polarized. Angles were determined using ImageJ Fiji software. For mouse cells, a total of 100 cells from each of 4 independent experiments were analyzed. For both HUVEC and MEC experiments, at least 60 cells from each of 3 independent experiments were analyzed. Statistics were computed using GraphPad Prism 7 and significance was evaluated using one-way ANOVA with Tukey post hoc test.

### Virus Production

For adenovirus virus production, VE-cad constructs were subcloned between *Bam*HI and *Age*I restriction sites in Gateway TagRFP-AS-N (Evrogen, Farmingdale, NY), in-frame with monomeric C-terminal RFP, then shuttled into pAd/Cmv/V5-DEST using LR Clonase recombination (Life Technologies). The vector was linearized using PacI and transfected into virus-producing QBI-293A cells. After 48-72 hours, cells were lysed and virus was harvested. To create replication-deficient second-generation lentivirus packaged with the indicated VE-cad gene containing a monomeric C-terminal RFP tag, the gene was cloned into pLenti6/V5-DEST and transfected into HEK-293T cells together with the necessary lentiviral regulatory genes. Lentivirus was collected from culture supernatants 48–72 hours after transfection and concentrated.

## Acknowledgements

We would like to thank Drs. B. Petrich and P. Vincent for reviewing the manuscript, members of the Kowalczyk laboratory for their help and advice and M. Myers for fruitful discussions. This work was supported by grants from the National Institutes of Health (R01AR050501 and R01AR048266 to A.P.K and HL095070 to KKG). This research project was supported in part by the Emory University Integrated Cellular Imaging Microscopy Core of the Winship Cancer Institute comprehensive cancer center grant, P30CA138292. This study was supported in part by the Mouse Transgenic and Gene Targeting Core (TMF) and the Emory Integrated Genomics Core (EIGC), which are subsidized by the Emory University School of Medicine and are Emory Integrated Core Facilities. Additional support was provided by the Georgia Clinical & Translational Science Alliance of the National Institutes of Health under Award Number UL1TR002378. The content is solely the responsibility of the authors and does not necessarily reflect the official views of the National Institutes of Health. The authors declare no competing financial interests.

## Author contributions

CGM, CMC, RHI, JC, KRM, TS, and MSH performed the experiments and analyzed results. WG performed data analysis. CGM, CMC, MSH, KRM, KKG, and APK designed the experiments. CGM and APK wrote the manuscript. All authors reviewed and approved the final version of the manuscript.

## Abbreviations

CBD: catenin binding domain
E: embryonic day
gRNA: guide RNA
HUVEC: human umbilical vein endothelial cells
JAIL: junction associated intermediate lamellipodia
JMD: juxtamembrane domain
LPS: lipopolysaccharide
MEC: microvascular endothelial cells
NHEJ: non-homologous end joining
P: postnatal day
p120: p120-catenin
DPBS: Dulbecco’s phosphate buffered saline
RFLP: restriction fragment length polymorphism
VE-cad: vascular endothelial cadherin

