## Supplemental Figures for "VE-cadherin endocytosis controls vascular integrity and patterning during development"

**A.**

**Number of postnatal progeny from cross:**
**VE-cad$^{GGG/GGG}$ X VE-Cad$^{STOP/+}$**

| Genotype | Expected | Observed |
| --- | --- | --- |
| *VE-cad$^{GGG/+}$* | 26 (50%) | 52 (100%) |
| *VE-cad$^{GGG/STOP}$* | 26 (50%) | ***0 (0%) |
| Total | | 52 |

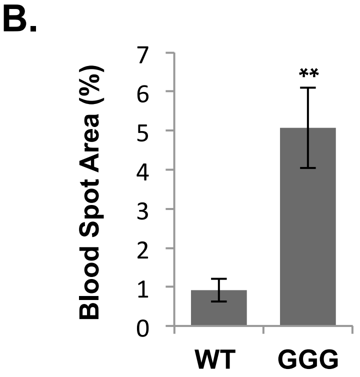


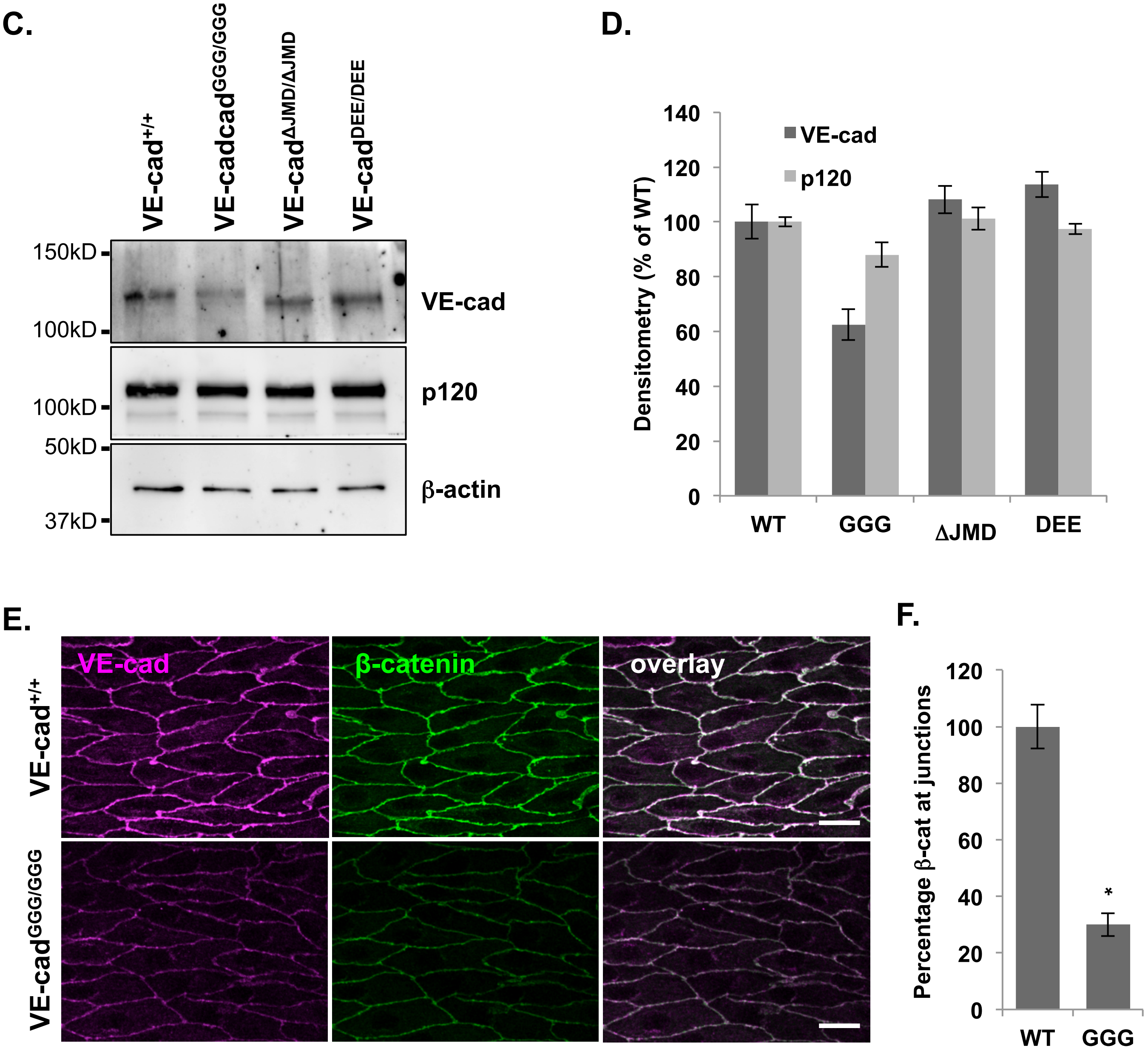


### Supplementary Figure 1

**Supplementary Figure 1**

**A,** Enhancement of the lethality phenotype in GGG mutants with one copy of VE-cad null VE-cad$^{STOP}$ allele. Genotyping analysis of offspring from VE-cad$^{GGG/GGG}$ × VE-cad$^{STOP/+}$ matings revealed no surviving VE-cad$^{GGG/STOP}$ pups. The expected number of mice was based on the total number of mice and expected Mendelian ratios. ***p<0.001 in $X^2$ analysis. **B,** Quantitation of blood spot area in VE-cad$^{GGG/GGG}$ mutant and wild type littermate retinas at P3. Area was quantitated from n= 7 (wild type) and n= 17 (VE-cad$^{GGG/GGG}$). *p<0.005. **C,** Isolated dermal endothelial cell lysates were prepared from the indicated WT and mutant mice and run in duplicate on SDS-PAGE gels. Protein expression levels were analyzed by western blotting using the indicated antibodies. **D,** Densitometry of western blot data, normalized to β-actin. Data was averaged from n=2 independent experiments and is presented as mean ± SEM. **E,** Decreased β-catenin at cell-cell borders in VE-cad$^{GGG/GGG}$ mutant mice. Aorta en face preparations were immunostained for VE-cad (red) and β-catenin (green). **F,** Quantitation of β-catenin levels at cell-cell junctions in the aortas of VE-cad$^{+/+}$ and VE-cad$^{GGG/GGG}$ mutant mice. β-catenin levels were decreased in the mutant similarly to VE-cad. Levels were quantitated from four independent experiments with 4-6 images per animal. Results represent the relative mean ± SEM. *p<0.05.

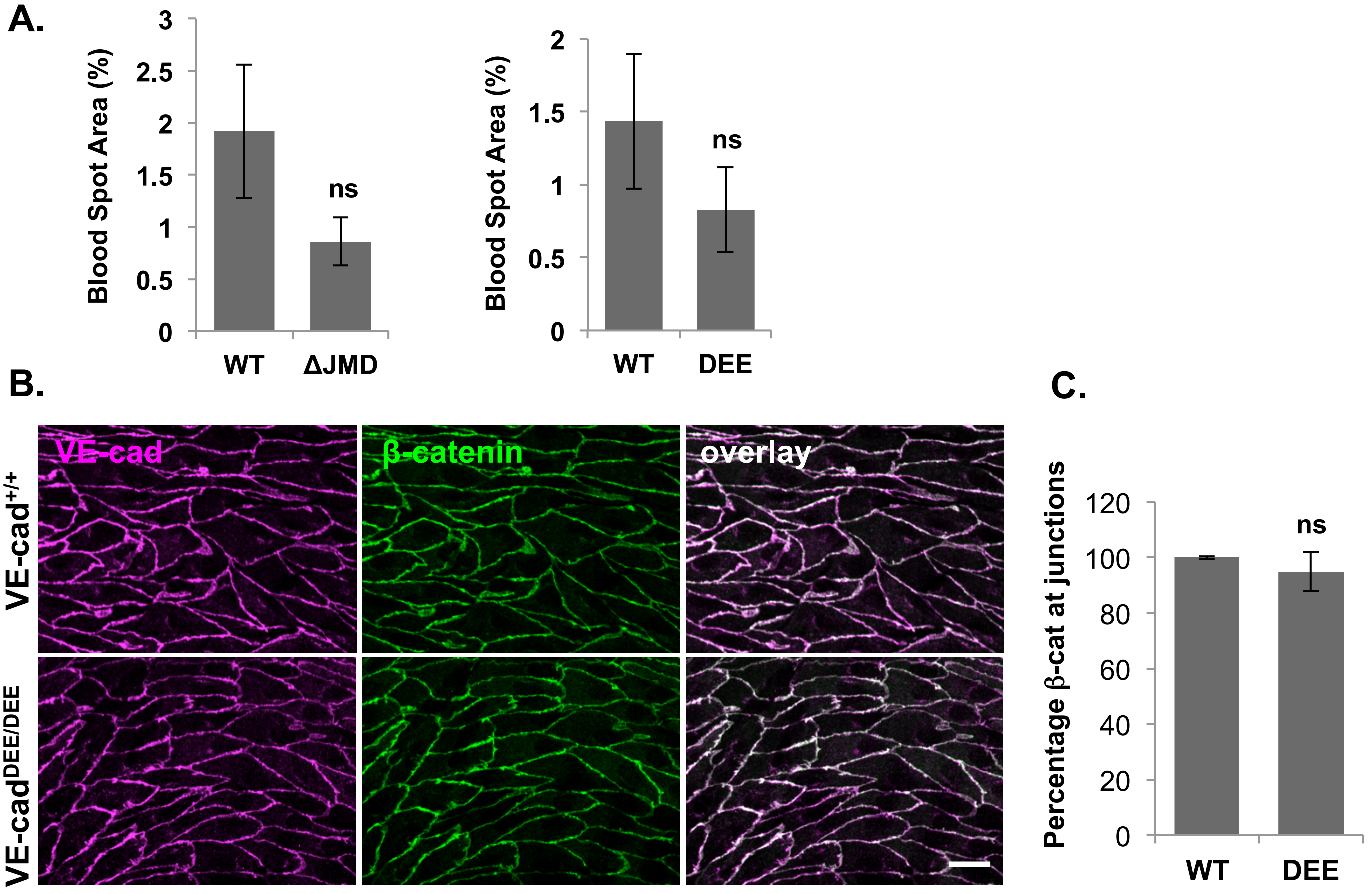


### Supplementary Figure 2

**Supplementary Figure 2**

**A,** Quantitation of blood spot area in the retinas of wild type and VE-cad $^{\Delta JMD/\Delta JMD}$ mutant littermates (left) and wild type and VE-cad$^{DEE/DEE}$ mutant littermates (right) at P3. Area was quantitated from n= 14 (wild type littermates of VE-cad $^{\Delta JMD/\Delta JMD}$) and n= 21 (VE-cad $^{\Delta JMD/\Delta JMD}$) and n= 12 (wild type littermates of VE-cad$^{DEE/DEE}$) and n= 16 (VE-cad$^{DEE/DEE}$); ns, not significant. **B,** Immunostaining analysis for VE-cad (red) and β-catenin (green) on *en face* aorta preparations from control and VE-cad $^{DEE/DEE}$ mutant mice. β-catenin levels and localization appeared normal in VE-cad $^{DEE/DEE}$ mutant mice. Scale bar: 20 μm. **C,** Quantitation of VE-cad and β-catenin levels at cell-cell junctions in the aortas of VE-cad$^{+/+}$ and VE-cad$^{DEE/DEE}$ mice. Levels were quantitated from three independent experiments with 4-6 images per animal, and shown as the relative mean ± SEM; ns, not significant.

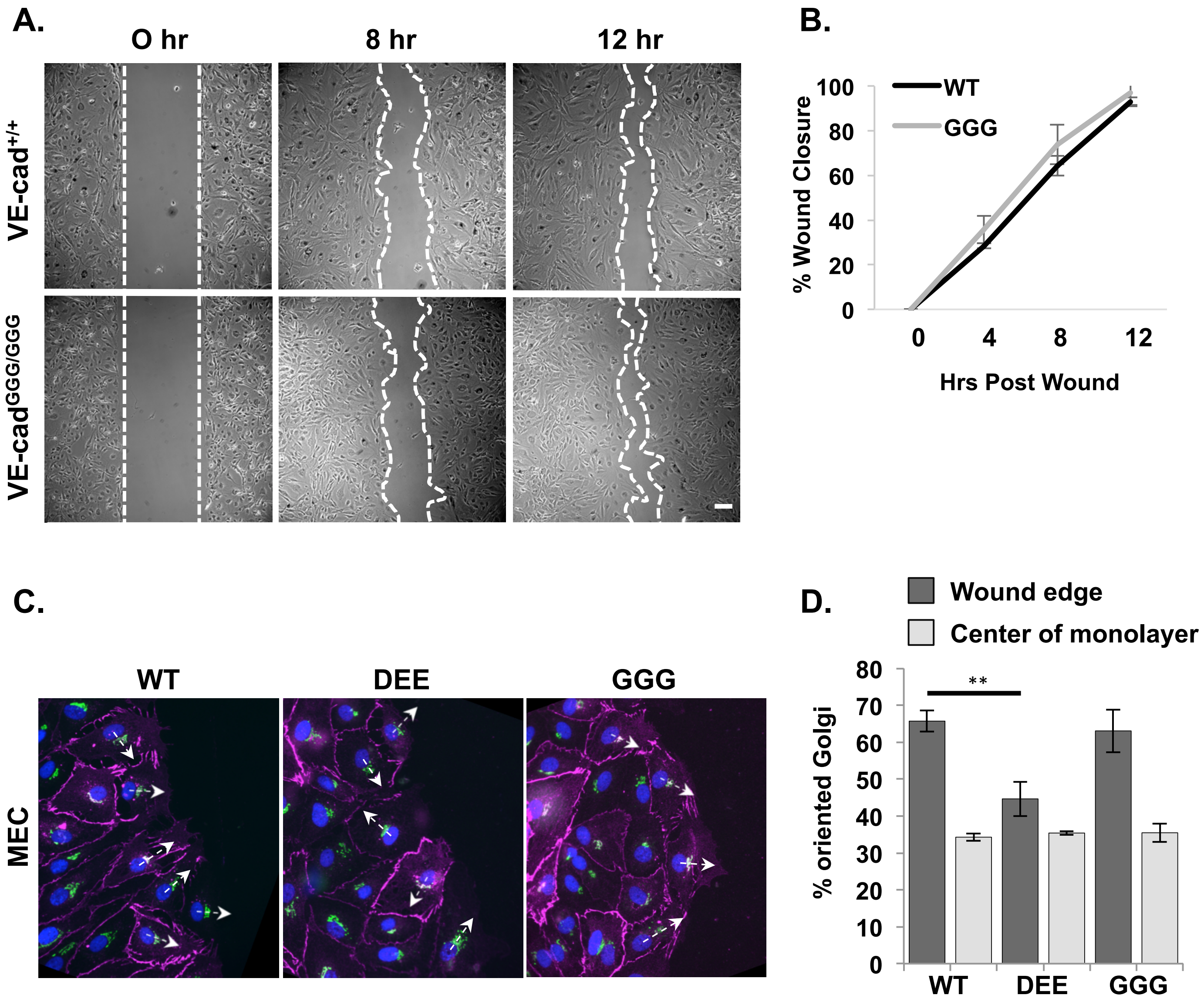


### Supplementary Figure 3

**Supplementary Figure 3**

**A,** Normal migration of isolated VE-cad$^{GGG/GGG}$ mutant dermal endothelial cells *in vitro*. Scratch-wound assays were performed with primary dermal microvascular endothelial cells isolated from VE-cad$^{+/+}$ and VE-cad$^{GGG/GGG}$ mutant mice. White dashed lines denote scratch borders. **B,** The percentage wound closure by VE-cad$^{+/+}$ and VE-cad$^{GGG/GGG}$ mutant cells was calculated over 12 hours using phase-contrast microscopy. Graph is representative of 2 independent experiments. **C,** Golgi reorientation defects in human microvascular endothelial cells expressing VE-cadDEE mutant. Human microvascular endothelial cells were adenovirally transduced with the indicated RFP tagged VE-cad constructs. Cells were fixed 6 hours after wounding and immunostained for RFP (red), GM-130 (green) and nuclei (blue) to visualize reorientation of the Golgi towards the wound edge. Arrows indicate the nucleus-Golgi polarity axis. **D,** Graph shows the average percentage ±SEM of RFP-positive cells with a polarized Golgi apparatus (in a 90° quadrant towards the wound edge) 6 hours after wounding. Graph represents average of 3 independent experiments. $^{**}p<0.005$.
